# Reemergence of cholera in Haiti

**DOI:** 10.1101/2022.10.31.514591

**Authors:** Daniel H. F. Rubin, Franz G. Zingl, Deborah R. Leitner, Ralph Ternier, Valusnor Compere, Samson Marseille, Damien Slater, Jason B. Harris, Fahima Chowdhury, Firdausi Qadri, Jacques Boncy, Louise C. Ivers, Matthew K. Waldor

## Abstract

Cholera was absent from Haiti until an inadvertent introduction by United Nations security forces in October 2010. The ensuing epidemic sickened 820,000 and caused 9,792 reported deaths^1^. The last cholera case in Haiti was recorded in January 2019, and in February 2022, Haiti was declared to have eliminated cholera^2^. In late September of 2022, a new outbreak began in Port-au-Prince and rapidly expanded to 964 suspected cases by mid-October of which 115 were confirmed by culture.^3^ Here, we present genomic and phenotypic analysis of the *Vibrio cholerae* isolated from a stool sample collected on September 30th, 2022 of an index case – a 10-year-old girl who presented with watery diarrhea and severe dehydration – to address the origins of the epidemic.

---

The 2022 *V. cholerae* isolate shares phenotypes with the 2010 outbreak strain. Both strains are *V. cholerae* serogroup O1 of the Ogawa serotype and have similar antibiograms, including resistance to trimethoprim/sulfamethoxazole and low-level resistance to ciprofloxacin (Supp. Table 1,2). This resistance profile is consistent among several other isolates from the current outbreak, suggesting that the strain isolated from the index case is representative of the ongoing epidemic.

To decipher the relationship between the current outbreak strain and other toxigenic O1 El Tor strains from the ongoing seventh pandemic of cholera, we sequenced the 9/30/2022 isolate, along with four 2021-2022 isolates from Dhaka, Bangladesh (Supp. Table 2). Phylogenetic analyses of over 1,200 isolates revealed that the 2022 Haiti index case clusters extremely closely with isolates from the 2010 outbreak in Haiti (Figure 1). This isolate is similarly closely related to 2010 Nepal isolates that were the origin of the initial outbreak as well as to 2013 strains from Mexico that were thought to have spread from Haiti^4^ and is divergent from currently circulating Bangladesh isolates. Haiti 2022 and Haiti 2010 isolates have identical *ctxB* (*ctxB7*) and other virulence factors (Supp. Table 3) and produce similar quantities of cholera toxin (Supp. Figure 1).

**Figure 1.**
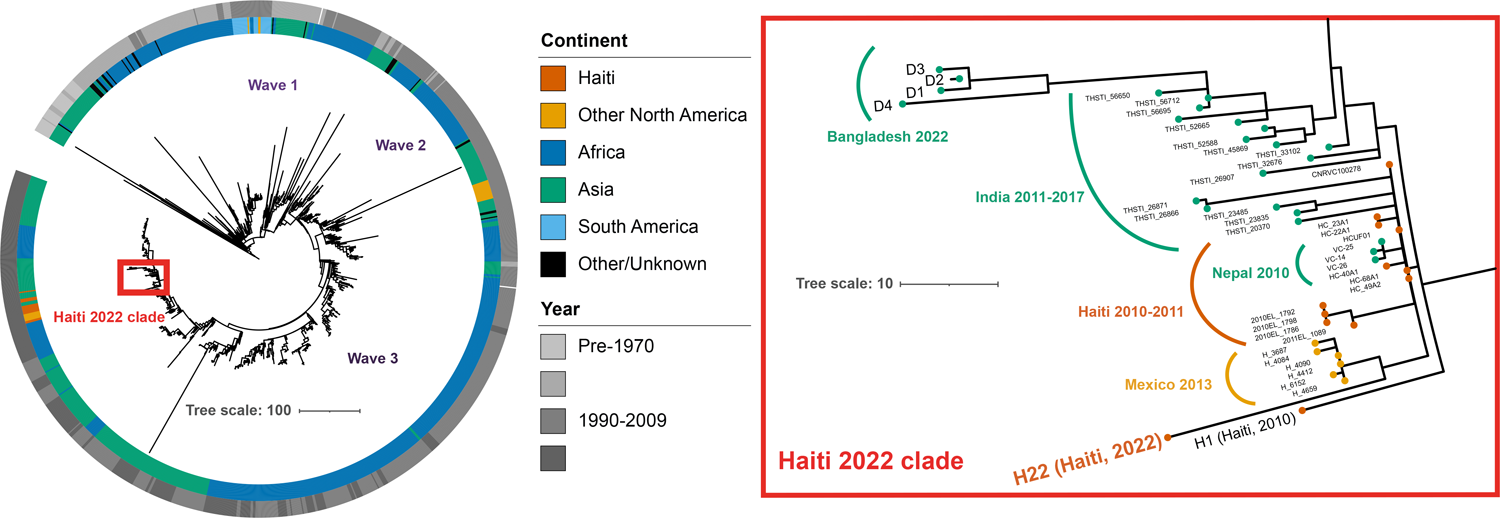
Phylogenetic tree of 7^th^ pandemic *Vibrio cholerae*. (Left) A phylogenetic tree of non-recombinogenic regions from 1,227 strains of O1 El Tor 7^th^ pandemic *V. cholerae*. Wave 1, wave 2, and wave 3 represent the dissemination of *V. cholerae* from Asia^6^. Tracks represent continent (inner) and year (outer) of isolation. Tree scale represents single-nucleotide polymorphisms (SNPs) per genome. (Right) Inset focused on the Haiti clade along with recent Asian isolates.

These analyses strongly suggest the reemergence of cholera in Haiti is caused by a descendant of the *V. cholerae* strain that gave rise to the 2010 epidemic. However, no cases of cholera were confirmed between 2019 and 2022 despite ongoing surveillance. Several explanations for the recrudescence of this strain are possible. The first is that toxigenic *V. cholerae* O1 persisted in Haiti through sub-clinical human infection and has recurred in the setting of waning population immunity coupled with a crisis in lack of clean water and sanitation. A second non-exclusive possibility is that this *V. cholerae* strain has persisted in environmental reservoirs. Finally, since the Haiti outbreak was ultimately transmitted to other countries in Latin America^4^, a third much less likely explanation given the absence of recent cholera cases in the region is that the current strain could have been reintroduced to Haiti from a nearby country. These findings, along with the resurgence of cholera in several parts of the world^5^ despite available tools, suggest that cholera control and prevention efforts must be redoubled.

## Acknowledgements

This work is funded by NIH R01AI-04237 and HHMI (MKW), NIH F30AI160911-01 and NIH T32GM007753 (DHFR), NIH R01HD102540 (JBH), and NIH R01AI099243 (LCI).

This article is subject to HHMI’s Open Access to Publications policy. HHMI lab heads have previously granted a nonexclusive CC BY 4.0 license to the public and a sublicensable license to HHMI in their research articles. Pursuant to those licenses, the author-accepted manuscript of this article can be made freely available under a CC BY 4.0 license immediately upon publication.

