## Supplemental Material for "Reemergence of cholera in Haiti"

Supplementary Appendix
Materials and Methods
Supplementary Tables 1-4
Supplementary Figure 1
References

### Methods and Materials

*Strains and Growth Conditions*

Strain H22 was imported from Haiti with CDC PHS Permit No. 20221004-3616A. Clinical samples of *Vibrio cholerae* (VC) were streaked overnight on lysogeny broth (LB) agar (Difco) plates. Individual colonies were then subcultured into LB (Difco) overnight. For induction of cholera toxin, strains were grown in AKI medium (1.5% Bacto peptone, 0.5% sodium chloride, 0.4% yeast extract, 0.3% sodium bicarbonate) with four hours stationary followed by four hours of shaking[1]. All growth was performed at 37°C. Clinical isolates presented in this study are presented in Supplementary Table 1.

*Cholera toxin analysis and serotyping*

Immunoblot analysis was performed as previously described[2]. Proteins were separated by SDS-PAGE using 4 to 12% NuPAGE Bis-Tris precast gels (Life Technologies) and transferred to nitrocellulose using an iBlot gel transfer device (Life Technologies). The prestained protein marker, SeeBlue (Invitrogen) was used as a molecular mass standard. Rabbit anti-CT polyclonal antibody (Abcam; catalog no. ab123129) and horseradish peroxidase (HRP)-linked anti-rabbit IgG were used as primary and secondary antibodies, respectively. Blots were developed with the SuperSignal West Pico Plus chemiluminescence substrate (Thermo Fisher) for five minutes and exposed in a ChemiDoc system (Bio-Rad Laboratories). This experiment was performed with three independent replicates. Serotyping was performed using slide agglutination with Ogawa- and Inaba-specific antisera (BD Difco).

*Minimum inhibitory concentration (MIC)-assays*

For MIC assays, overnight cultures grown in LB were shifted into fresh LB at a final OD600 of 0.02. Three-fold serial dilutions of antimicrobial agents were tested across the following range of concentrations: 100µg/ml to 0.000564503µg/ml. After 18 h incubation of growth at 37°C, $OD_{600}$ was measured. The MIC was defined as the minimum antimicrobial agent concentration which inhibited bacterial growth. Growth was defined by an at least four-fold increase of the $OD_{600}$ compared to a sterile control. All MICs were performed with four independent replicates.

Whole-genome Sequencing and Phylogenetic Analysis

Genomic DNA (gDNA) was first isolated from *V. cholerae* strains using the GeneJet genomic DNA purification kit (Thermo Fisher). DNA was fragmented using a sonicator (Peak power 50, duty factor 10, cycles per burst 200, duration 90 seconds) (Covaris M220). Fragments were then prepared for short read sequencing via the NEBNext Ultra II DNA Library Prep Kit (NEB). Libraries were then sequenced on a NextSeq 550 (Illumina).

For phylogenetic analysis, genomes[2-21] (Supp. Table 4) were analyzed as previously published[2,22]. Raw reads were downloaded from the European Nucleotide Archive. For fully assembled genomes, pseudoreads of 100-bp overlapping reads were generated using either wgsim (https://github.com/lh3/wgsim) or fasta-to-fastq.pl (https://github.com/ekg/fasta-to-fastq/blob/master/fasta_to_fastq.pl). Reads were then mapped to the genome of N16961 (GenBank accession nos. LT907989 and LT907990), a seventh pandemic wave 1 isolate, with Snippy v4.6.0 (https://github.com/tseemann/snippy) using freebayes v1.3.6 (https://github.com/freebayes/freebayes) with a minimum read coverage of 4, minimum base quality of 13, a mapping quality of 60, and a requirement of 75% read concordance. Alignments for previously published isolates were kindly provided previously by D. Domman (University of New Mexico) and alignments from Mexican isolates can be found in previously published work[22]. The known repeat region TLC-RS1-CTX was masked from the genome, as was the recombinogenic VSP-II region using Geneious (Biomatters). Recombinogenic sites were then further masked with Gubbins v3.3[23], and a phylogenetic tree was assembled based on the remaining 8,191 polymorphisms with RAxML v8.212[24] using a general time reversible with gamma rate heterogeneity (GTRGAMMA) model and 100 bootstraps. The resulting tree was visualized in iTOL v6[25].

Contigs were assembled using SPAdes v3.15.5[26] and annotated using Geneious (Biomatters) from the H1 genome (aka KW3, Genbank accession no. GCA_001318185.1). Alleles of interest were identified through either direct mapping of reads to the H1 or N16961 genomes via Geneious or through BLAST[27] of the N16961 allele on assembled genomes.

#### Data Availability

All accession numbers for isolates will be available publicly prior to publication.

**Supplementary Table 1**
**Representative antibiogram of *V. cholerae* isolates from the Laboratoire National de Santé Publique (LNSP).**

| **Antibiotic** | MIC (μg/mL) | Interpretation | Kirby-Bauer (Interpretation) |
|---|---|---|---|
| Ampicillin 10 mcg | 16 | I | |
| Trimethoprim/Sulfamethoxazole | ≥320 | R | |
| Tetracycline | | | S |
| Cefoxitin | 16 | I | |
| Ceftazidime | 0.5 | S | |
| Imipenem | 2 | S | |
| Piperacillin/Tazobactam | ≤4 | S | |
| Cefuroxime | ≤1 | | |
| Azithromycin | 0.4 | | S |

**Supplementary Table 2**
**Isolates sequenced in this study and strain characteristics.**

| Metadata | D1 | D2 | D3 | D4 | H22 | H1 |
|---|---|---|---|---|---|---|
| **Date of isolation** | 10/31/2021 | 01/12/2022 | 03/23/2022 | 03/30/2022 | 09/30/2022 | 2010 |
| **Location of isolation** | Dhaka, Bangladesh | Dhaka, Bangladesh | Dhaka, Bangladesh | Dhaka, Bangladesh | Port-au-Prince, Haiti | Haiti |
| **Serotype** | Inaba | Inaba | Inaba | Ogawa | Ogawa | Ogawa |

| MIC | | | | | | |
|---|---|---|---|---|---|---|
| Ciprofloxacin | 0.4 | 0.4 | 0.4 | 0.4 | 0.4 | 0.4 |
| Sulfamethoxazole | >100 | >100 | >100 | >100 | >100 | >100 |
| Trimethoprim | 100 | >100 | >100 | 100 | >100 | >100 |
| Doxycycline | 0.1 | 0.1 | 0.1 | 0.1 | 0.1 | 0.1 |
| Ceftriaxone | 0.02 | 0.02 | 0.02 | 0.03 | 0.02 | 0.02 |
| Tetracycline | 0.4 | 0.4 | 0.4 | 0.4 | 0.4 | 0.4 |
| Erythromycin | 4 | 4 | 4 | 4 | 4 | 4 |
| Trimethoprim: Sulfamethoxazole 1:5 | >100 | >100 | >100 | >100 | >100 | >100 |
| Azithromycin | 0.4 | 0.4 | 0.4 | 0.4 | 0.4 | 0.4 |
| Gentamicin | 11 | 11 | 11 | 11 | 33 | 33 |

**Supplementary Table 3**
**Pathogenicity factors from strains in our study.** Changes to alleles are called with reference to the N16961 genome.

| Gene | D1 | D2 | D3 | D4 | H22 | H1 |
|---|---|---|---|---|---|---|
| *wbeT* | Tn insertion (c.324) | Tn insertion (c.324) | Tn insertion (c.324) | Intact | Intact | Intact |
| *ctxB* | ctxB7 | ctxB7 | ctxB7 | ctxB7 | ctxB7 | ctxB7 |
| *tcpA* | c.A266>G | c.A266>G | c.A266>G | c.A266>G | c.A266>G | c.A266>G |
| *vspII* | Del VC_0495-VC_0512::ISVch4) | Del VC_0495-VC_0512::ISVch4) | Del VC_0495-VC_0512::ISVch4) | Del VC_0495-VC_0512::ISVch4) | Del VC_0495-VC_0512::ISVch4) | Del VC_0495-VC_0512::ISVch4) |

**Supplementary Figure 1**

**Cholera toxin (CT) western blot analysis**. Shown is a representative immunoblot loaded with equal amounts of sterile-filtered supernatants of AKI-grown D1 (lane1), D2 (lane 2), D3 (lane 3), D4 (lane 4), H22 (lane 5) and H1 (lane 6). The sizes of the A and B subunit (A and B) of CT are indicated by arrows on the right. Lines to the left indicate the molecular masses of the protein standards in kDa. The gap in between D4 and H22 contained samples not analyzed for this study.

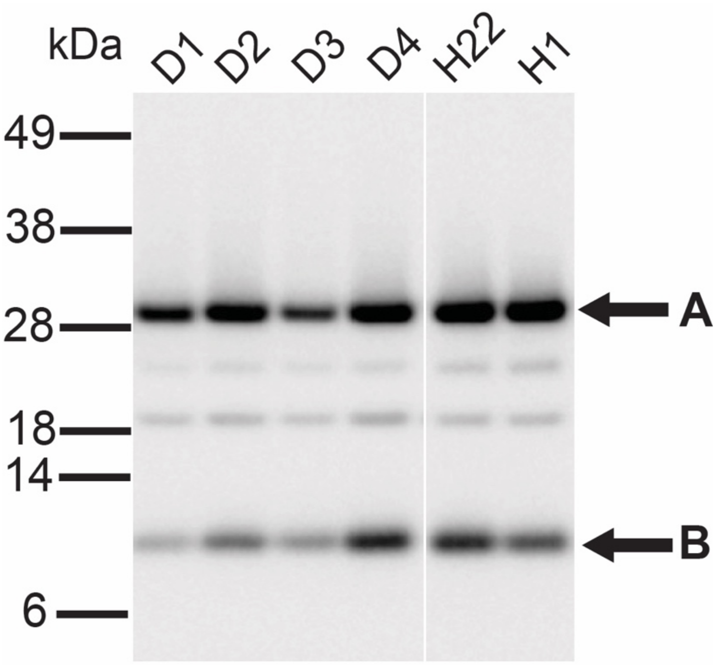

**Supplementary Table 4**
**Toxigenic O1 *Vibrio cholerae* genomes and accessions used in Figure 1.**

| Isolate name | Year | Country | Continent | EBI-ENA accession no. | Sequence ID | GenBank accession no. | Source |
|---|---|---|---|---|---|---|---|
| **7994** | 2009 | India | Asia | ERR351210 | ERR351210 | | Abd El Ghany et al. PLoS Negl Trop Dis 2014 |
| **7772** | 2009 | India | Asia | ERR351208 | ERR351208 | | Abd El Ghany et al. PLoS Negl Trop Dis 2014 |
| **7683** | 2009 | India | Asia | ERR351207 | ERR351207 | | Abd El Ghany et al. PLoS Negl Trop Dis 2014 |
| **6801** | 2009 | India | Asia | ERR351206 | ERR351206 | | Abd El Ghany et al. PLoS Negl Trop Dis 2014 |
| **6734** | 2009 | India | Asia | ERR351205 | ERR351205 | | Abd El Ghany et al. PLoS Negl Trop Dis 2014 |
| **6557** | 2009 | India | Asia | ERR351203 | ERR351203 | | Abd El Ghany et al. PLoS Negl Trop Dis 2014 |
| **6259** | 2009 | India | Asia | ERR351199 | ERR351199 | | Abd El Ghany et al. PLoS Negl Trop Dis 2014 |
| **6016** | 2009 | India | Asia | ERR351193 | ERR351193 | | Abd El Ghany et al. PLoS Negl Trop Dis 2014 |
| **5663** | 2009 | India | Asia | ERR351192 | ERR351192 | | Abd El Ghany et al. PLoS Negl Trop Dis 2014 |
| **5417** | 2009 | India | Asia | ERR351191 | ERR351191 | | Abd El Ghany et al. PLoS Negl Trop Dis 2014 |
| **5286** | 2009 | India | Asia | ERR351188 | ERR351188 | | Abd El Ghany et al. PLoS Negl Trop Dis 2014 |
| **5202** | 2009 | India | Asia | ERR351185 | ERR351185 | | Abd El Ghany et al. PLoS Negl Trop Dis 2014 |
| **5185** | 2009 | India | Asia | ERR351179 | ERR351179 | | Abd El Ghany et al. PLoS Negl Trop Dis 2014 |
| **5046** | 2009 | India | Asia | ERR351175 | ERR351175 | | Abd El Ghany et al. PLoS Negl Trop Dis 2014 |
| **4966** | 2009 | India | Asia | ERR351174 | ERR351174 | | Abd El Ghany et al. PLoS Negl Trop Dis 2014 |
| **UG026** | 2015 | Uganda | Africa | | UG026 | SAMN08744332 | Bwire et al. PLoS Negl Trop Dis 2018 |
| **UG010** | 2016 | Uganda | Africa | | UG010 | SAMN08744330 | Bwire et al. PLoS Negl Trop Dis 2018 |
| **UG020** | 2016 | Uganda | Africa | | UG020 | SAMN08744331 | Bwire et al. PLoS Negl Trop Dis 2018 |
| **H1** | 2010 | Haiti | North America | | H1 | GCA_000275645.1 | Chin et al. NEJM 2011 |
| **MJ1236** | 1994 | Bangladesh | Asia | | MJ1236 | CP001485/CP001486 | Chun et al. Proc Natl Acad Sci USA 2009 |
| **CIRS101** | 2002 | Bangladesh | Asia | | CIRS101 | ACVW00000000 | Chun et al. Proc Natl Acad Sci USA 2009 |
| **MO10** | 1992 | India | Asia | | MO10 | AAKF03000000 | Chun et al. Proc Natl Acad Sci USA 2009 |

| | | | | | | | |
|---|---|---|---|---|---|---|---|
| **RC9** | 1985 | Kenya | Africa | | RC9 | ACHX00000000 | Chun et al. Proc Natl Acad Sci USA 2009 |
| **B33** | 2004 | Mozambique | Africa | | B33 | ACHZ00000000 | Chun et al. Proc Natl Acad Sci USA 2009 |
| **V01-CDC** | 1991 | Chile | South America | ERR576950 | ERR576950 | | Didelot et al. PLoS Genet 2015 |
| **1525-CDC** | 1994 | China | Asia | ERR579986 | ERR579986 | | Didelot et al. PLoS Genet 2015 |
| **2578-CDC** | 1985 | China | Asia | ERR579919 | ERR579919 | | Didelot et al. PLoS Genet 2015 |
| **200106-CDC** | 2001 | China | Asia | ERR579313 | ERR579313 | | Didelot et al. PLoS Genet 2015 |
| **129-CDC** | 1994 | China | Asia | ERR579311 | ERR579311 | | Didelot et al. PLoS Genet 2015 |
| **9-CDC** | 2001 | China | Asia | ERR579309 | ERR579309 | | Didelot et al. PLoS Genet 2015 |
| **SC1998_098-1-CDC** | 1998 | China | Asia | ERR579114 | ERR579114 | | Didelot et al. PLoS Genet 2015 |
| **JX1998-1306-1-CDC** | 1998 | China | Asia | ERR579093 | ERR579093 | | Didelot et al. PLoS Genet 2015 |
| **SC1998_242-1-CDC** | 1998 | China | Asia | ERR579092 | ERR579092 | | Didelot et al. PLoS Genet 2015 |
| **SC2000-CDC** | 2000 | China | Asia | ERR579091 | ERR579091 | | Didelot et al. PLoS Genet 2015 |
| **VC2248-1-CDC** | 2008 | China | Asia | ERR579090 | ERR579090 | | Didelot et al. PLoS Genet 2015 |
| **VC2454-CDC** | 2005 | China | Asia | ERR579089 | ERR579089 | | Didelot et al. PLoS Genet 2015 |
| **1283-CDC** | 1984 | China | Asia | ERR579088 | ERR579088 | | Didelot et al. PLoS Genet 2015 |
| **132-CDC** | 1978 | China | Asia | ERR579087 | ERR579087 | | Didelot et al. PLoS Genet 2015 |
| **143-CDC** | 2001 | China | Asia | ERR579086 | ERR579086 | | Didelot et al. PLoS Genet 2015 |
| **147-CDC** | 2002 | China | Asia | ERR579085 | ERR579085 | | Didelot et al. PLoS Genet 2015 |
| **1551-CDC** | 1979 | China | Asia | ERR579083 | ERR579083 | | Didelot et al. PLoS Genet 2015 |
| **1575-CDC** | 1990 | China | Asia | ERR579082 | ERR579082 | | Didelot et al. PLoS Genet 2015 |
| **1583-CDC** | 1991 | China | Asia | ERR579081 | ERR579081 | | Didelot et al. PLoS Genet 2015 |
| **1593-CDC** | 1992 | China | Asia | ERR579080 | ERR579080 | | Didelot et al. PLoS Genet 2015 |
| **1605-CDC** | 1993 | China | Asia | ERR579079 | ERR579079 | | Didelot et al. PLoS Genet 2015 |
| **1627-CDC** | 1997 | China | Asia | ERR579078 | ERR579078 | | Didelot et al. PLoS Genet 2015 |
| **1909-CDC** | 1998 | China | Asia | ERR579077 | ERR579077 | | Didelot et al. PLoS Genet 2015 |

| | | | | | | | |
|---|---|---|---|---|---|---|---|
| **1944-CDC** | 2000 | China | Asia | ERR579076 | ERR579076 | | Didelot et al. PLoS Genet 2015 |
| **2255-CDC** | 2008 | China | Asia | ERR579075 | ERR579075 | | Didelot et al. PLoS Genet 2015 |
| **2308-CDC** | 1978 | China | Asia | ERR579074 | ERR579074 | | Didelot et al. PLoS Genet 2015 |
| **2530-CDC** | 1996 | China | Asia | ERR579073 | ERR579073 | | Didelot et al. PLoS Genet 2015 |
| **2574-CDC** | 1980 | China | Asia | ERR579072 | ERR579072 | | Didelot et al. PLoS Genet 2015 |
| **2580-CDC** | 1989 | China | Asia | ERR579070 | ERR579070 | | Didelot et al. PLoS Genet 2015 |
| **2605-CDC** | 1998 | China | Asia | ERR579069 | ERR579069 | | Didelot et al. PLoS Genet 2015 |
| **2657-CDC** | 1995 | China | Asia | ERR579068 | ERR579068 | | Didelot et al. PLoS Genet 2015 |
| **2710-CDC** | 1986 | China | Asia | ERR579067 | ERR579067 | | Didelot et al. PLoS Genet 2015 |
| **2744-CDC** | 1962 | China | Asia | ERR579066 | ERR579066 | | Didelot et al. PLoS Genet 2015 |
| **2752-CDC** | 1981 | China | Asia | ERR579065 | ERR579065 | | Didelot et al. PLoS Genet 2015 |
| **2757-CDC** | 1987 | China | Asia | ERR579064 | ERR579064 | | Didelot et al. PLoS Genet 2015 |
| **2783-CDC** | 1964 | China | Asia | ERR579063 | ERR579063 | | Didelot et al. PLoS Genet 2015 |
| **2833-CDC** | 1979 | China | Asia | ERR579062 | ERR579062 | | Didelot et al. PLoS Genet 2015 |
| **2848-CDC** | 1988 | China | Asia | ERR579061 | ERR579061 | | Didelot et al. PLoS Genet 2015 |
| **2865-CDC** | 1961 | China | Asia | ERR579060 | ERR579060 | | Didelot et al. PLoS Genet 2015 |
| **2873-CDC** | 1961 | China | Asia | ERR579059 | ERR579059 | | Didelot et al. PLoS Genet 2015 |
| **2881-CDC** | 1961 | China | Asia | ERR579058 | ERR579058 | | Didelot et al. PLoS Genet 2015 |
| **2894-CDC** | 1961 | China | Asia | ERR579057 | ERR579057 | | Didelot et al. PLoS Genet 2015 |
| **2914-CDC** | 1966 | China | Asia | ERR579056 | ERR579056 | | Didelot et al. PLoS Genet 2015 |
| **2921-CDC** | 1962 | China | Asia | ERR579055 | ERR579055 | | Didelot et al. PLoS Genet 2015 |
| **2957-CDC** | 1964 | China | Asia | ERR579054 | ERR579054 | | Didelot et al. PLoS Genet 2015 |
| **2981-CDC** | 1965 | China | Asia | ERR579053 | ERR579053 | | Didelot et al. PLoS Genet 2015 |
| **3014-CDC** | 1969 | China | Asia | ERR579052 | ERR579052 | | Didelot et al. PLoS Genet 2015 |
| **3024-CDC** | 1973 | China | Asia | ERR579051 | ERR579051 | | Didelot et al. PLoS Genet 2015 |

| | | | | | | | |
|---|---|---|---|---|---|---|---|
| **3075-CDC** | 1974 | China | Asia | ERR579050 | ERR579050 | | Didelot et al. PLoS Genet 2015 |
| **3270-CDC** | 1977 | China | Asia | ERR579049 | ERR579049 | | Didelot et al. PLoS Genet 2015 |
| **3289-CDC** | 1978 | China | Asia | ERR577143 | ERR577143 | | Didelot et al. PLoS Genet 2015 |
| **3304-CDC** | 1979 | China | Asia | ERR577142 | ERR577142 | | Didelot et al. PLoS Genet 2015 |
| **3497-CDC** | 1980 | China | Asia | ERR577141 | ERR577141 | | Didelot et al. PLoS Genet 2015 |
| **3538-CDC** | 1981 | China | Asia | ERR577140 | ERR577140 | | Didelot et al. PLoS Genet 2015 |
| **3564-CDC** | 1982 | China | Asia | ERR577139 | ERR577139 | | Didelot et al. PLoS Genet 2015 |
| **3595-CDC** | 1983 | China | Asia | ERR576996 | ERR576996 | | Didelot et al. PLoS Genet 2015 |
| **3626-CDC** | 1983 | China | Asia | ERR576995 | ERR576995 | | Didelot et al. PLoS Genet 2015 |
| **3633-CDC** | 1984 | China | Asia | ERR576988 | ERR576988 | | Didelot et al. PLoS Genet 2015 |
| **3735-CDC** | 1988 | China | Asia | ERR576987 | ERR576987 | | Didelot et al. PLoS Genet 2015 |
| **4039-CDC** | 1993 | China | Asia | ERR576986 | ERR576986 | | Didelot et al. PLoS Genet 2015 |
| **4070-CDC** | 1994 | China | Asia | ERR576985 | ERR576985 | | Didelot et al. PLoS Genet 2015 |
| **4210-CDC** | 1999 | China | Asia | ERR576984 | ERR576984 | | Didelot et al. PLoS Genet 2015 |
| **63244-CDC** | 1963 | China | Asia | ERR576980 | ERR576980 | | Didelot et al. PLoS Genet 2015 |
| **642345-CDC** | 1964 | China | Asia | ERR576979 | ERR576979 | | Didelot et al. PLoS Genet 2015 |
| **93284-CDC** | 1993 | China | Asia | ERR576977 | ERR576977 | | Didelot et al. PLoS Genet 2015 |
| **936-CDC** | 1998 | China | Asia | ERR576976 | ERR576976 | | Didelot et al. PLoS Genet 2015 |
| **956-CDC** | 2001 | China | Asia | ERR576975 | ERR576975 | | Didelot et al. PLoS Genet 2015 |
| **AHV1003-CDC** | 2010 | China | Asia | ERR576974 | ERR576974 | | Didelot et al. PLoS Genet 2015 |
| **D118-CDC** | 1961 | China | Asia | ERR576972 | ERR576972 | | Didelot et al. PLoS Genet 2015 |
| **GD196110-CDC** | 1961 | China | Asia | ERR576971 | ERR576971 | | Didelot et al. PLoS Genet 2015 |
| **V011506-CDC** | 2001 | China | Asia | ERR576951 | ERR576951 | | Didelot et al. PLoS Genet 2015 |
| **6310-CDC** | 1961 | Indonesia | Asia | ERR576983 | ERR576983 | | Didelot et al. PLoS Genet 2015 |
| **6311-CDC** | 1961 | Indonesia | Asia | ERR576982 | ERR576982 | | Didelot et al. PLoS Genet 2015 |

| | | | | | | | |
|---|---|---|---|---|---|---|---|
| **6312-CDC** | 1961 | Indonesia | Asia | ERR576981 | ERR576981 | | Didelot et al. PLoS Genet 2015 |
| **863-CDC** | 1986 | Mauritania | Africa | ERR576978 | ERR576978 | | Didelot et al. PLoS Genet 2015 |
| **X190-CDC** | 1991 | Peru | South America | ERR580006 | ERR580006 | | Didelot et al. PLoS Genet 2015 |
| **C6706-CDC** | 1991 | Peru | South America | ERR576973 | ERR576973 | | Didelot et al. PLoS Genet 2015 |
| **T21-CDC** | 1990 | Thailand | Asia | ERR576970 | ERR576970 | | Didelot et al. PLoS Genet 2015 |
| **CNRVC14012 6** | 1991 | Bolivia | South America | ERR976497 | 16244_7_46 | | Domman et al. Science 2017 |
| **CNRVC14012 5** | 1992 | Bolivia | South America | ERR976496 | 16244_7_45 | | Domman et al. Science 2017 |
| **CNRVC14012 4** | 1992 | Bolivia | South America | ERR976495 | 16244_7_44 | | Domman et al. Science 2017 |
| **CNRVC14012 3** | 1992 | Bolivia | South America | ERR976494 | 16244_7_43 | | Domman et al. Science 2017 |
| **CNRVC14011 7** | 1991 | Bolivia | South America | ERR976488 | 16244_7_37 | | Domman et al. Science 2017 |
| **CNRVC14012 2** | 1992 | Brazil | South America | ERR976493 | 16244_7_42 | | Domman et al. Science 2017 |
| **CNRVC14012 1** | 1991 | Brazil | South America | ERR976492 | 16244_7_41 | | Domman et al. Science 2017 |
| **CNRVC14011 8** | 1991 | Brazil | South America | ERR976489 | 16244_7_38 | | Domman et al. Science 2017 |
| **CNRVC95001 4** | 1995 | Ecuador | South America | ERR1879579 | CNRVC950014_ATGTCA_L0 02 | | Domman et al. Science 2017 |
| **CNRVC94018 3** | 1994 | French Guiana | South America | ERR1879578 | CNRVC940183_AGTTCC_L0 02 | | Domman et al. Science 2017 |
| **CNRVC93002 5** | 1993 | French Guiana | South America | ERR1879554 | CNRVC930025_TCCCGA_L0 01 | | Domman et al. Science 2017 |
| **54267_1994** | 1994 | Mexico | North America | ERR163241 | 8014_8_9 | | Domman et al. Science 2017 |
| **33297_1993** | 1993 | Mexico | North America | ERR163240 | 8014_8_8 | | Domman et al. Science 2017 |
| **60483_1995** | 1995 | Mexico | North America | ERR163244 | 8014_8_12 | | Domman et al. Science 2017 |
| **60452_1995** | 1995 | Mexico | North America | ERR163243 | 8014_8_11 | | Domman et al. Science 2017 |
| **54328_1994** | 1994 | Mexico | North America | ERR163242 | 8014_8_10 | | Domman et al. Science 2017 |
| **85** | 2000 | Mexico | North America | ERR108522 | 7138_2_7 | | Domman et al. Science 2017 |
| **54** | 1999 | Mexico | North America | ERR108518 | 7138_2_3 | | Domman et al. Science 2017 |
| **688** | 2006 | Mexico | North America | ERR108537 | 7138_2_22 | | Domman et al. Science 2017 |
| **838** | 1999 | Mexico | North America | ERR108517 | 7138_2_2 | | Domman et al. Science 2017 |

| | | | | | | | |
|---|---|---|---|---|---|---|---|
| **82** | 1998 | Mexico | North America | ERR108516 | 7138_2_1 | | Domman et al. Science 2017 |
| **87662** | 1993 | Mexico | North America | ERR044786 | 6437_7_9 | | Domman et al. Science 2017 |
| **87667** | 1993 | Mexico | North America | ERR044785 | 6437_7_8 | | Domman et al. Science 2017 |
| **116075** | 1992 | Mexico | North America | ERR044783 | 6437_7_6 | | Domman et al. Science 2017 |
| **116073** | 1991 | Mexico | North America | ERR044781 | 6437_7_4 | | Domman et al. Science 2017 |
| **116072** | 1991 | Mexico | North America | ERR044779 | 6437_7_2 | | Domman et al. Science 2017 |
| **95430** | 1997 | Mexico | North America | ERR044791 | 6437_7_14 | | Domman et al. Science 2017 |
| **Mex6** | 1992 | Mexico | North America | ERR042754 | 6353_8_3 | | Domman et al. Science 2017 |
| **H_3687** | 2013 | Mexico | North America | ERR466829 | 12005_1_11 | | Domman et al., Science 2017 |
| **H_4084** | 2013 | Mexico | North America | ERR466834 | 12005_1_16 | | Domman et al., Science 2017 |
| **H_4090** | 2013 | Mexico | North America | ERR466835 | 12005_1_17 | | Domman et al., Science 2017 |
| **H_4412** | 2013 | Mexico | North America | ERR466846 | 12014_1_28 | | Domman et al., Science 2017 |
| **H_4659** | 2013 | Mexico | North America | ERR471121 | 12022_1_32 | | Domman et al., Science 2017 |
| **H_6152** | 2013 | Mexico | North America | ERR471128 | 12022_1_39 | | Domman et al., Science 2017 |
| **ZChol** | 2016 | Zambia | Africa | SRR17283076/SRR17283077 | ZChol | | Fakoya et al. mBio 2022 |
| **S152** | 2003 | Mozambique | Africa | ERS1938085 | 152_L008_R1 | | Garrine et al. PLoS Neglect Trop Dis 2017 |
| **S121** | 2003 | Mozambique | Africa | ERS1938079 | 121_L008_R1 | | Garrine et al. PLoS Neglect Trop Dis 2017 |
| **S1020229_6** | 2010 | Mozambique | Africa | ERS1938072 | 1020229_6_L007_R1 | | Garrine et al. PLoS Neglect Trop Dis 2017 |
| **S1019828_5** | 2010 | Mozambique | Africa | ERS1938068 | 1019828_5_L007_R1 | | Garrine et al. PLoS Neglect Trop Dis 2017 |
| **S0074** | 2003 | Mozambique | Africa | ERS1938060 | 0074_L002_R1 | | Garrine et al. PLoS Neglect Trop Dis 2017 |
| **S0035** | 2002 | Mozambique | Africa | ERS1938059 | 0035_L008_R1 | | Garrine et al. PLoS Neglect Trop Dis 2017 |
| **S0034** | 2002 | Mozambique | Africa | ERS1938058 | 0034_L008_R1 | | Garrine et al. PLoS Neglect Trop Dis 2017 |
| **S0014** | 2002 | Mozambique | Africa | ERS1938051 | 0014_L008_R1 | | Garrine et al. PLoS Neglect Trop Dis 2017 |
| **HC_23A1** | 2010 | Haiti | North America | SRR135546 | SRR135546 | | Hasan et al. PNAS 2012 |
| **HC_49A2** | 2010 | Haiti | North America | SRR135544 | SRR135544 | | Hasan et al. PNAS 2012 |

| | | | | | | | |
|---|---|---|---|---|---|---|---|
| **N16961** | 1975 | Bangladesh | Asia | | Vibrio_cholerae_O1_biovar_el tor_str_N16961_v2 | LT907989/LT907990 | Heidelberg et al. Nature 2000 |
| **VC-26** | 2010 | Nepal | Asia | SRR308727 | SRR308727 | | Hendriksen et al. Mbio 2011 |
| **VC-26** | 2010 | Nepal | Asia | SRR308726 | SRR308726 | | Hendriksen et al. Mbio 2011 |
| **VC-22** | 2010 | Nepal | Asia | SRR308725 | SRR308725 | | Hendriksen et al. Mbio 2011 |
| **VC-21** | 2010 | Nepal | Asia | SRR308724 | SRR308724 | | Hendriksen et al. Mbio 2011 |
| **VC-20** | 2010 | Nepal | Asia | SRR308723 | SRR308723 | | Hendriksen et al. Mbio 2011 |
| **VC-19** | 2010 | Nepal | Asia | SRR308722 | SRR308722 | | Hendriksen et al. Mbio 2011 |
| **VC-18** | 2010 | Nepal | Asia | SRR308721 | SRR308721 | | Hendriksen et al. Mbio 2011 |
| **VC-17** | 2010 | Nepal | Asia | SRR308720 | SRR308720 | | Hendriksen et al. Mbio 2011 |
| **VC-16** | 2010 | Nepal | Asia | SRR308717 | SRR308717 | | Hendriksen et al. Mbio 2011 |
| **VC-15** | 2010 | Nepal | Asia | SRR308716 | SRR308716 | | Hendriksen et al. Mbio 2011 |
| **VC-14** | 2010 | Nepal | Asia | SRR308715 | SRR308715 | | Hendriksen et al. Mbio 2011 |
| **VC-13** | 2010 | Nepal | Asia | SRR308713 | SRR308713 | | Hendriksen et al. Mbio 2011 |
| **VC-12** | 2010 | Nepal | Asia | SRR308709 | SRR308709 | | Hendriksen et al. Mbio 2011 |
| **VC-11** | 2010 | Nepal | Asia | SRR308708 | SRR308708 | | Hendriksen et al. Mbio 2011 |
| **VC-10** | 2010 | Nepal | Asia | SRR308707 | SRR308707 | | Hendriksen et al. Mbio 2011 |
| **VC-9** | 2010 | Nepal | Asia | SRR308706 | SRR308706 | | Hendriksen et al. Mbio 2011 |
| **VC-8** | 2010 | Nepal | Asia | SRR308705 | SRR308705 | | Hendriksen et al. Mbio 2011 |
| **VC-7** | 2010 | Nepal | Asia | SRR308704 | SRR308704 | | Hendriksen et al. Mbio 2011 |
| **VC-6** | 2010 | Nepal | Asia | SRR308703 | SRR308703 | | Hendriksen et al. Mbio 2011 |
| **VC-5** | 2010 | Nepal | Asia | SRR308693 | SRR308693 | | Hendriksen et al. Mbio 2011 |
| **VC-4** | 2010 | Nepal | Asia | SRR308692 | SRR308692 | | Hendriksen et al. Mbio 2011 |
| **VC-3** | 2010 | Nepal | Asia | SRR308691 | SRR308691 | | Hendriksen et al. Mbio 2011 |
| **VC-2** | 2010 | Nepal | Asia | SRR308690 | SRR308690 | | Hendriksen et al. Mbio 2011 |
| **VC-1** | 2010 | Nepal | Asia | SRR308665 | SRR308665 | | Hendriksen et al. Mbio 2011 |

| | | | | | | | |
|---|---|---|---|---|---|---|---|
| **Tanz_100_1** | 2015 | Tanzania | Africa | ERS2318718 | IL100065177_S148_L008 | | Kachwamba et al. BMC Infect Dis 2017 |
| **Tanz_78** | 2015 | Tanzania | Africa | ERS2318712 | IL100065171_S142_L008 | | Kachwamba et al. BMC Infect Dis 2017 |
| **Tanz_65** | 2015 | Tanzania | Africa | ERS2318708 | IL100065167_S138_L008 | | Kachwamba et al. BMC Infect Dis 2017 |
| **Tanz_60** | 2012 | Tanzania | Africa | ERS2318705 | IL100065164_S135_L008 | | Kachwamba et al. BMC Infect Dis 2017 |
| **Tanz_58** | 2011 | Tanzania | Africa | ERS2318704 | IL100065163_S134_L008 | | Kachwamba et al. BMC Infect Dis 2017 |
| **Tanz_56** | 2012 | Tanzania | Africa | ERS2318703 | IL100065162_S133_L008 | | Kachwamba et al. BMC Infect Dis 2017 |
| **Tanz_48** | 2015 | Tanzania | Africa | ERS2318700 | IL100064947_S130_L008 | | Kachwamba et al. BMC Infect Dis 2017 |
| **Tanz_42** | 2015 | Tanzania | Africa | ERS2318697 | IL100064944_S127_L008 | | Kachwamba et al. BMC Infect Dis 2017 |
| **Tanz_33** | 2015 | Tanzania | Africa | ERS2318693 | IL100064940_S123_L008 | | Kachwamba et al. BMC Infect Dis 2017 |
| **Tanz_20** | 2015 | Tanzania | Africa | ERS2318689 | IL100064936_S119_L008 | | Kachwamba et al. BMC Infect Dis 2017 |
| **Tanz_14** | 2015 | Tanzania | Africa | ERS2318685 | IL100064932_S115_L008 | | Kachwamba et al. BMC Infect Dis 2017 |
| **Tanz_9** | 2015 | Tanzania | Africa | ERS2318682 | IL100064929_S45_L002 | | Kachwamba et al. BMC Infect Dis 2017 |
| **Tanz_3** | 2015 | Tanzania | Africa | ERS2318681 | IL100064928_S44_L002 | | Kachwamba et al. BMC Infect Dis 2017 |
| **Tanz_2** | 2015 | Tanzania | Africa | ERS2318680 | IL100064927_S43_L002 | | Kachwamba et al. BMC Infect Dis 2017 |
| **CP1050** | 2010 | Bangladesh | Asia | SRR227335 | SRR227335 | | Katz et al. Mbio 2013 |
| **CP1048** | 2010 | Bangladesh | Asia | SRR227303 | SRR227303 | | Katz et al. Mbio 2013 |
| **2010EL_1749** | 2010 | Cameroon | Africa | SRR773655 | 2010EL_1749 | | Katz et al. Mbio 2013 |
| **HC-22A1** | 2010 | Haiti | North America | SRR191381 | SRR191381 | | Katz et al. Mbio 2013 |
| **HC-68A1** | 2010 | Haiti | North America | SRR191363 | SRR191363 | | Katz et al. Mbio 2013 |
| **HC-40A1** | 2010 | Haiti | North America | SRR135545 | SRR135545 | | Katz et al. Mbio 2013 |
| **HCUF01** | 2010 | Haiti | North America | SRR135540 | SRR135540 | | Katz et al. Mbio 2013 |
| **2011EL_1137** | 2009 | Republic of South Africa | Africa | SRR774784 | 2011EL_1137 | | Katz et al. Mbio 2013 |
| **CP1042** | 2010 | Thailand | Asia | SRR227324 | SRR227324 | | Katz et al. Mbio 2013 |
| **CP1041** | 2004 | Zambia | Africa | SRR227309 | SRR227309 | | Katz et al. Mbio 2013 |
| **CP1038** | 2009 | Zimbabwe | Africa | SRR227311 | SRR227311 | | Katz et al. Mbio 2013 |
| **KNC1709** | 2009 | Kenya | Africa | ERR114418 | 7346_8_53 | | Kiiru et al. Plos One 2013 |

| | | | | | | | |
|---|---|---|---|---|---|---|---|
| **KNC8678** | 2009 | Kenya | Africa | ERR114416 | 7346_8_51 | | Kiiru et al. Plos One 2013 |
| **KNC8673** | 2009 | Kenya | Africa | ERR114415 | 7346_8_50 | | Kiiru et al. Plos One 2013 |
| **KNC151** | 2010 | Kenya | Africa | ERR117573 | 7298_7_7 | | Kiiru et al. Plos One 2013 |
| **KNC135** | 2009 | Kenya | Africa | ERR117572 | 7298_7_6 | | Kiiru et al. Plos One 2013 |
| **KNC145** | 2010 | Kenya | Africa | ERR117571 | 7298_7_5 | | Kiiru et al. Plos One 2013 |
| **KNC1509** | 2009 | Kenya | Africa | ERR117613 | 7298_7_47 | | Kiiru et al. Plos One 2013 |
| **KNC8572** | 2009 | Kenya | Africa | ERR117612 | 7298_7_46 | | Kiiru et al. Plos One 2013 |
| **KNC8675** | 2009 | Kenya | Africa | ERR117610 | 7298_7_44 | | Kiiru et al. Plos One 2013 |
| **KNC207** | 2009 | Kenya | Africa | ERR117609 | 7298_7_43 | | Kiiru et al. Plos One 2013 |
| **KNC1420** | 2009 | Kenya | Africa | ERR117607 | 7298_7_41 | | Kiiru et al. Plos One 2013 |
| **KNC8679** | 2008 | Kenya | Africa | ERR117606 | 7298_7_40 | | Kiiru et al. Plos One 2013 |
| **KNC233** | 2009 | Kenya | Africa | ERR117605 | 7298_7_39 | | Kiiru et al. Plos One 2013 |
| **KNC206** | 2009 | Kenya | Africa | ERR117604 | 7298_7_38 | | Kiiru et al. Plos One 2013 |
| **KNC231** | 2009 | Kenya | Africa | ERR117600 | 7298_7_34 | | Kiiru et al. Plos One 2013 |
| **KNC8669** | 2010 | Kenya | Africa | ERR117599 | 7298_7_33 | | Kiiru et al. Plos One 2013 |
| **KNC155** | 2010 | Kenya | Africa | ERR117597 | 7298_7_31 | | Kiiru et al. Plos One 2013 |
| **KNE168** | 2010 | Kenya | Africa | ERR117569 | 7298_7_3 | | Kiiru et al. Plos One 2013 |
| **KNC8885** | 2010 | Kenya | Africa | ERR117595 | 7298_7_29 | | Kiiru et al. Plos One 2013 |
| **KNC156** | 2009 | Kenya | Africa | ERR117594 | 7298_7_28 | | Kiiru et al. Plos One 2013 |
| **KNC1888** | 2007 | Kenya | Africa | ERR117593 | 7298_7_27 | | Kiiru et al. Plos One 2013 |
| **KNC157** | 2009 | Kenya | Africa | ERR117592 | 7298_7_26 | | Kiiru et al. Plos One 2013 |
| **KNC161** | 2010 | Kenya | Africa | ERR117590 | 7298_7_24 | | Kiiru et al. Plos One 2013 |
| **KNC147** | 2010 | Kenya | Africa | ERR117588 | 7298_7_22 | | Kiiru et al. Plos One 2013 |
| **KNC11** | 2010 | Kenya | Africa | ERR117586 | 7298_7_20 | | Kiiru et al. Plos One 2013 |
| **KNC158** | 2010 | Kenya | Africa | ERR117583 | 7298_7_17 | | Kiiru et al. Plos One 2013 |
| **KNC8889** | 2010 | Kenya | Africa | ERR117581 | 7298_7_15 | | Kiiru et al. Plos One 2013 |
| **KNC133** | 2007 | Kenya | Africa | ERR117579 | 7298_7_13 | | Kiiru et al. Plos One 2013 |
| **KNC8880** | 2010 | Kenya | Africa | ERR117578 | 7298_7_12 | | Kiiru et al. Plos One 2013 |

| | | | | | | | |
|---|---|---|---|---|---|---|---|
| **KNC56** | 2010 | Kenya | Africa | ERR117577 | 7298_7_11 | | Kiiru et al. Plos One 2013 |
| **KNEXXH** | 2009 | Kenya | Africa | ERR117556 | 7298_6_86 | | Kiiru et al. Plos One 2013 |
| **KNEXC** | 2009 | Kenya | Africa | ERR117555 | 7298_6_85 | | Kiiru et al. Plos One 2013 |
| **KNE3C** | 2010 | Kenya | Africa | ERR117473 | 7298_5_3 | | Kiiru et al. Plos One 2013 |
| **KNE3G** | 2010 | Kenya | Africa | ERR117490 | 7298_5_20 | | Kiiru et al. Plos One 2013 |
| **KNE170** | 2010 | Kenya | Africa | ERR117485 | 7298_5_15 | | Kiiru et al. Plos One 2013 |
| **KNE11B** | 2010 | Kenya | Africa | ERR117484 | 7298_5_14 | | Kiiru et al. Plos One 2013 |
| **KNE134B** | 2009 | Kenya | Africa | ERR117482 | 7298_5_12 | | Kiiru et al. Plos One 2013 |
| **KNE134** | 2009 | Kenya | Africa | ERR117480 | 7298_5_10 | | Kiiru et al. Plos One 2013 |
| **VE3** | 2010 | Kenya | Africa | ERR037741 | 6084_8_20 | | Kiiru et al. Plos One 2013 |
| **VE2** | 2010 | Kenya | Africa | ERR037739 | 6084_8_19 | | Kiiru et al. Plos One 2013 |
| **VE1** | 2010 | Kenya | Africa | ERR037738 | 6084_8_18 | | Kiiru et al. Plos One 2013 |
| **YA00122542** | 2018 | Zimbabwe | Africa | ERR3342516 | YA00122542-VCH_S14_L001 | | Mashe et al. NEJM 2020 |
| **YA00122540** | 2018 | Zimbabwe | Africa | ERR3342515 | YA00122540-VCH_S12_L001 | | Mashe et al. NEJM 2020 |
| **YA00122539** | 2018 | Zimbabwe | Africa | ERR3342514 | YA00122539-VCH_S11_L001 | | Mashe et al. NEJM 2020 |
| **YA00122538** | 2018 | Zimbabwe | Africa | ERR3342513 | YA00122538-VCH_S23_L001 | | Mashe et al. NEJM 2020 |
| **YA00122536** | 2018 | Zimbabwe | Africa | ERR3342512 | YA00122536-VCH_S22_L001 | | Mashe et al. NEJM 2020 |
| **YA00122534** | 2018 | Zimbabwe | Africa | ERR3342511 | YA00122534-VCH_S20_L001 | | Mashe et al. NEJM 2020 |
| **YA00122533** | 2018 | Zimbabwe | Africa | ERR3342510 | YA00122533-VCH_S19_L001 | | Mashe et al. NEJM 2020 |
| **YA00122532** | 2018 | Zimbabwe | Africa | ERR3342509 | YA00122532-VCH_S18_L001 | | Mashe et al. NEJM 2020 |
| **YA00122531** | 2018 | Zimbabwe | Africa | ERR3342508 | YA00122531-VCH_S17_L001 | | Mashe et al. NEJM 2020 |
| **YA00122530** | 2018 | Zimbabwe | Africa | ERR3342507 | YA00122530-VCH_S16_L001 | | Mashe et al. NEJM 2020 |
| **YA00120881** | 2018 | Zimbabwe | Africa | ERR3342506 | YA00120881-VCH_S10_L001 | | Mashe et al. NEJM 2020 |
| **A5** | 1989 | Angola | Africa | ERR025381 | 5174_7_4 | | Mutreja et al. Nature 2011 |
| **A316** | 1993 | Argentina | South America | ERR018181 | 4056_8_8 | | Mutreja et al. Nature 2011 |
| **A200** | 1992 | Argentina | South America | ERR018168 | 4056_7_8 | | Mutreja et al. Nature 2011 |
| **A201** | 1992 | Argentina | South America | ERR018167 | 4056_7_7 | | Mutreja et al. Nature 2011 |
| **A186** | 1992 | Argentina | South America | ERR018158 | 4056_7_1 | | Mutreja et al. Nature 2011 |

| | | | | | | | |
|---|---|---|---|---|---|---|---|
| **GP143** | 1978 | Bahrein | Asia | ERR018192 | 4075_3_6 | | Mutreja et al. Nature 2011 |
| **MG116226** | 1991 | Bangladesh | Asia | ERR025396 | 5174_8_9 | | Mutreja et al. Nature 2011 |
| **A22** | 1979 | Bangladesh | Asia | ERR025386 | 5174_7_9 | | Mutreja et al. Nature 2011 |
| **A19** | 1971 | Bangladesh | Asia | ERR025385 | 5174_7_8 | | Mutreja et al. Nature 2011 |
| **A10** | 1979 | Bangladesh | Asia | ERR025383 | 5174_7_6 | | Mutreja et al. Nature 2011 |
| **4662** | 2001 | Bangladesh | Asia | ERR025373 | 5174_6_7 | | Mutreja et al. Nature 2011 |
| **A487(2)** | 2007 | Bangladesh | Asia | ERR018182 | 4056_8_9 | | Mutreja et al. Nature 2011 |
| **A346(2)** | 1994 | Bangladesh | Asia | ERR018179 | 4056_8_6 | | Mutreja et al. Nature 2011 |
| **A390** | 1987 | Bangladesh | Asia | ERR018177 | 4056_8_4 | | Mutreja et al. Nature 2011 |
| **A397** | 1987 | Bangladesh | Asia | ERR018176 | 4056_8_3 | | Mutreja et al. Nature 2011 |
| **A488(2)** | 2006 | Bangladesh | Asia | ERR018128 | 4056_2_7 | | Mutreja et al. Nature 2011 |
| **MG116025** | 1991 | Bangladesh | Asia | ERR018122 | 4056_2_12 | | Mutreja et al. Nature 2011 |
| **A383** | 2002 | Bangladesh | Asia | ERR018121 | 4056_2_11 | | Mutreja et al. Nature 2011 |
| **MJ1485** | 1994 | Bangladesh | Asia | ERR018120 | 4056_2_10 | | Mutreja et al. Nature 2011 |
| **4672** | 2000 | Bangladesh | Asia | ERR019884 | 3002_8_6 | | Mutreja et al. Nature 2011 |
| **4670** | 1991 | Bangladesh | Asia | ERR019883 | 3002_8_5 | | Mutreja et al. Nature 2011 |
| **4661** | 2001 | Bangladesh | Asia | ERR018116 | 2956_6_6 | | Mutreja et al. Nature 2011 |
| **4663** | 2001 | Bangladesh | Asia | ERR018115 | 2956_6_5 | | Mutreja et al. Nature 2011 |
| **4679** | 1999 | Bangladesh | Asia | ERR018114 | 2956_6_4 | | Mutreja et al. Nature 2011 |
| **4675** | 2001 | Bangladesh | Asia | ERR018113 | 2956_6_3 | | Mutreja et al. Nature 2011 |
| **A193** | 1992 | Bolivia | South America | ERR018160 | 4056_7_11 | | Mutreja et al. Nature 2011 |
| **A185** | 1992 | Colombia | South America | ERR018156 | 4056_6_9 | | Mutreja et al. Nature 2011 |
| **A177** | 1992 | Colombia | South America | ERR018149 | 4056_6_2 | | Mutreja et al. Nature 2011 |
| **A180** | 1992 | Colombia | South America | ERR018148 | 4056_6_12 | | Mutreja et al. Nature 2011 |
| **A184** | 1992 | Colombia | South America | ERR018146 | 4056_6_10 | | Mutreja et al. Nature 2011 |
| **A481** | 2007 | Djibouti | Africa | ERR018175 | 4056_8_2 | | Mutreja et al. Nature 2011 |
| **A482** | 2007 | Djibouti | Africa | ERR018174 | 4056_8_12 | | Mutreja et al. Nature 2011 |
| **A483** | 2007 | Djibouti | Africa | ERR018172 | 4056_8_10 | | Mutreja et al. Nature 2011 |

| | | | | | | | |
|---|---|---|---|---|---|---|---|
| **A18** | 1977 | India | Asia | ERR025384 | 5174_7_7 | | Mutreja et al. Nature 2011 |
| **4536** | 2007 | India | Asia | ERR025375 | 5174_6_9 | | Mutreja et al. Nature 2011 |
| **4519** | 2005 | India | Asia | ERR025374 | 5174_6_8 | | Mutreja et al. Nature 2011 |
| **4646** | 2007 | India | Asia | ERR025372 | 5174_6_6 | | Mutreja et al. Nature 2011 |
| **4600** | 2007 | India | Asia | ERR025371 | 5174_6_5 | | Mutreja et al. Nature 2011 |
| **4488** | 2006 | India | Asia | ERR025369 | 5174_6_3 | | Mutreja et al. Nature 2011 |
| **4552** | 2007 | India | Asia | ERR025368 | 5174_6_2 | | Mutreja et al. Nature 2011 |
| **4585** | 2007 | India | Asia | ERR025366 | 5174_6_1 | | Mutreja et al. Nature 2011 |
| **4339** | 2004 | India | Asia | ERR025361 | 5174_5_6 | | Mutreja et al. Nature 2011 |
| **4538** | 2007 | India | Asia | ERR025360 | 5174_5_5 | | Mutreja et al. Nature 2011 |
| **4593** | 2007 | India | Asia | ERR025359 | 5174_5_4 | | Mutreja et al. Nature 2011 |
| **4623** | 2007 | India | Asia | ERR025358 | 5174_5_3 | | Mutreja et al. Nature 2011 |
| **4551** | 2007 | India | Asia | ERR025357 | 5174_5_2 | | Mutreja et al. Nature 2011 |
| **PRL64** | 1992 | India | Asia | ERR018195 | 4075_3_9 | | Mutreja et al. Nature 2011 |
| **PRL5** | 1980 | India | Asia | ERR018189 | 4075_3_3 | | Mutreja et al. Nature 2011 |
| **GP152** | 1979 | India | Asia | ERR018188 | 4075_3_2 | | Mutreja et al. Nature 2011 |
| **IDHO1'726** | 2009 | India | Asia | ERR018187 | 4075_3_12 | | Mutreja et al. Nature 2011 |
| **GP60** | 1973 | India | Asia | ERR018186 | 4075_3_11 | | Mutreja et al. Nature 2011 |
| **A130** | 1989 | India | Asia | ERR018154 | 4056_6_7 | | Mutreja et al. Nature 2011 |
| **A131** | 1989 | India | Asia | ERR018153 | 4056_6_6 | | Mutreja et al. Nature 2011 |
| **MBRN14** | 2004 | India | Asia | ERR018130 | 4056_2_9 | | Mutreja et al. Nature 2011 |
| **V5** | 1989 | India | Asia | ERR018127 | 4056_2_6 | | Mutreja et al. Nature 2011 |
| **V109** | 1990 | India | Asia | ERR018126 | 4056_2_5 | | Mutreja et al. Nature 2011 |
| **V212-1** | 1991 | India | Asia | ERR018125 | 4056_2_4 | | Mutreja et al. Nature 2011 |
| **VC51** | 1992 | India | Asia | ERR018124 | 4056_2_3 | | Mutreja et al. Nature 2011 |
| **MBN17** | 2004 | India | Asia | ERR018123 | 4056_2_2 | | Mutreja et al. Nature 2011 |
| **A330** | 1993 | India | Asia | ERR018119 | 4056_2_1 | | Mutreja et al. Nature 2011 |
| **4322** | 2004 | India | Asia | ERR019881 | 3002_8_3 | | Mutreja et al. Nature 2011 |

| | | | | | | | |
|---|---|---|---|---|---|---|---|
| **4656** | 2006 | India | Asia | ERR018112 | 2956_6_2 | | Mutreja et al. Nature 2011 |
| **4605** | 2007 | India | Asia | ERR018111 | 2956_6_1 | | Mutreja et al. Nature 2011 |
| **A6** | 1957 | Indonesia | Asia | ERR025382 | 5174_7_5 | | Mutreja et al. Nature 2011 |
| **7685** | 2009 | Kenya | Africa | ERR028076 | 4370_3_8 | | Mutreja et al. Nature 2011 |
| **7686** | 2009 | Kenya | Africa | ERR028075 | 4370_3_7 | | Mutreja et al. Nature 2011 |
| **7687** | 2009 | Kenya | Africa | ERR028074 | 4370_3_6 | | Mutreja et al. Nature 2011 |
| **7684** | 2009 | Kenya | Africa | ERR028068 | 4370_3_11 | | Mutreja et al. Nature 2011 |
| **7682** | 2009 | Kenya | Africa | ERR028066 | 4370_3_1 | | Mutreja et al. Nature 2011 |
| **6215** | 2005 | Kenya | Africa | ERR019297 | 4370_2_9 | | Mutreja et al. Nature 2011 |
| **6193** | 2005 | Kenya | Africa | ERR019296 | 4370_2_8 | | Mutreja et al. Nature 2011 |
| **6194** | 2007 | Kenya | Africa | ERR019295 | 4370_2_7 | | Mutreja et al. Nature 2011 |
| **6197** | 2007 | Kenya | Africa | ERR019292 | 4370_2_4 | | Mutreja et al. Nature 2011 |
| **6201** | 2007 | Kenya | Africa | ERR019291 | 4370_2_3 | | Mutreja et al. Nature 2011 |
| **6210** | 2007 | Kenya | Africa | ERR019290 | 4370_2_2 | | Mutreja et al. Nature 2011 |
| **6212** | 2007 | Kenya | Africa | ERR019289 | 4370_2_12 | | Mutreja et al. Nature 2011 |
| **6191** | 2005 | Kenya | Africa | ERR019288 | 4370_2_11 | | Mutreja et al. Nature 2011 |
| **6214** | 2007 | Kenya | Africa | ERR019287 | 4370_2_10 | | Mutreja et al. Nature 2011 |
| **GP140** | 1978 | Malaysia | Asia | ERR018193 | 4075_3_7 | | Mutreja et al. Nature 2011 |
| **A231** | 1991 | Mexico | North America | ERR018161 | 4056_7_12 | | Mutreja et al. Nature 2011 |
| **A232** | 1991 | Mexico | North America | ERR018159 | 4056_7_10 | | Mutreja et al. Nature 2011 |
| **1627** | 2005 | Mozambique | Africa | ERR025377 | 5174_7_1 | | Mutreja et al. Nature 2011 |
| **1362** | 2005 | Mozambique | Africa | ERR025367 | 5174_6_10 | | Mutreja et al. Nature 2011 |
| **1346** | 2005 | Mozambique | Africa | ERR025356 | 5174_5_1 | | Mutreja et al. Nature 2011 |
| **A152** | 1991 | Mozambique | Africa | ERR018152 | 4056_6_5 | | Mutreja et al. Nature 2011 |
| **A154** | 1991 | Mozambique | Africa | ERR018151 | 4056_6_4 | | Mutreja et al. Nature 2011 |
| **A155** | 1991 | Mozambique | Africa | ERR018150 | 4056_6_3 | | Mutreja et al. Nature 2011 |
| **A32** | 1991 | Peru | South America | ERR025390 | 5174_8_3 | | Mutreja et al. Nature 2011 |
| **A31** | 1991 | Peru | South America | ERR025389 | 5174_8_2 | | Mutreja et al. Nature 2011 |

| | | | | | | | |
|---|---|---|---|---|---|---|---|
| **A29** | 1991 | Peru | South America | ERR025388 | 5174_8_1 | | Mutreja et al. Nature 2011 |
| **A27** | 1991 | Peru | South America | ERR025378 | 5174_7_10 | | Mutreja et al. Nature 2011 |
| **4784** | 2009 | Tanzania | Africa | ERR025370 | 5174_6_4 | | Mutreja et al. Nature 2011 |
| **A4** | 1973 | Unknown | | ERR025380 | 5174_7_3 | | Mutreja et al. Nature 2011 |
| **A109** | 1990 | Unknown | | ERR018147 | 4056_6_11 | | Mutreja et al. Nature 2011 |
| **4113** | 2003 | Vietnam | Asia | ERR025364 | 5174_5_9 | | Mutreja et al. Nature 2011 |
| **4121** | 2004 | Vietnam | Asia | ERR025362 | 5174_5_7 | | Mutreja et al. Nature 2011 |
| **A245** | 1989 | Vietnam | Asia | ERR018171 | 4056_8_1 | | Mutreja et al. Nature 2011 |
| **A241** | 1989 | Vietnam | Asia | ERR018169 | 4056_7_9 | | Mutreja et al. Nature 2011 |
| **4122** | 2007 | Vietnam | Asia | ERR019885 | 3002_8_7 | | Mutreja et al. Nature 2011 |
| **4111** | 2002 | Vietnam | Asia | ERR019880 | 3002_8_2 | | Mutreja et al. Nature 2011 |
| **4110** | 1995 | Vietnam | Asia | ERR019879 | 3002_8_1 | | Mutreja et al. Nature 2011 |
| **2010EL_1798** | 2010 | Haiti | North America | | 2010EL_1798 | AELI00000000.1 | Reimer et al. EID 2011 |
| **2010EL_1792** | 2010 | Haiti | North America | | 2010EL_1792 | AELJ00000000.1 | Reimer et al. EID 2011 |
| **2010EL_1786** | 2010 | Haiti | North America | | 2010EL_1786 | CP003069.1/CP003070.1 | Reimer et al. EID 2011 |
| **2011EL_1089** | 2010 | Haiti | North America | SRR773660 | 2011EL_1089 | | Reimer et al. EID 2011 |
| **S9KCH9** | 2010 | Pakistan | Asia | ERR051772 | 6714_6_5 | | Shah et al. EID 2014 |
| **S7KCH20** | 2010 | Pakistan | Asia | ERR051770 | 6714_6_3 | | Shah et al. EID 2014 |
| **S4KCH16** | 2010 | Pakistan | Asia | ERR051767 | 6714_5_23 | | Shah et al. EID 2014 |
| **S2KCH17** | 2010 | Pakistan | Asia | ERR051765 | 6714_5_21 | | Shah et al. EID 2014 |
| **S23HH18** | 2010 | Pakistan | Asia | ERR051786 | 6714_6_19 | | Shah et al. EID 2014 |
| **S6KCH7** | 2010 | Pakistan | Asia | ERR051769 | 6714_6_2 | | Shah et al. EID 2014 |
| **S5KCH10** | 2010 | Pakistan | Asia | ERR051768 | 6714_6_1 | | Shah et al. EID 2014 |
| **F8D25** | 2010 | Pakistan | Asia | ERR051752 | 6714_5_8 | | Shah et al. EID 2014 |
| **S14P9** | 2010 | Pakistan | Asia | ERR051777 | 6714_6_10 | | Shah et al. EID 2014 |
| **F4D48** | 2010 | Pakistan | Asia | ERR051748 | 6714_5_4 | | Shah et al. EID 2014 |
| **S26R24** | 2010 | Pakistan | Asia | ERR051789 | 6714_6_22 | | Shah et al. EID 2014 |
| **F1DN4** | 2010 | Pakistan | Asia | ERR051745 | 6714_5_1 | | Shah et al. EID 2014 |

| | | | | | | | |
|---|---|---|---|---|---|---|---|
| **S27RG11** | 2010 | Pakistan | Asia | ERR051790 | 6714_6_23 | | Shah et al. EID 2014 |
| **S24RG6** | 2010 | Pakistan | Asia | ERR051787 | 6714_6_20 | | Shah et al. EID 2014 |
| **F14KPD3** | 2010 | Pakistan | Asia | ERR051758 | 6714_5_14 | | Shah et al. EID 2014 |
| **F2D59** | 2010 | Pakistan | Asia | ERR051746 | 6714_5_2 | | Shah et al. EID 2014 |
| **F11D4** | 2010 | Pakistan | Asia | ERR051755 | 6714_5_11 | | Shah et al. EID 2014 |
| **F7D30** | 2010 | Pakistan | Asia | ERR051751 | 6714_5_7 | | Shah et al. EID 2014 |
| **S22HH17** | 2010 | Pakistan | Asia | ERR051785 | 6714_6_18 | | Shah et al. EID 2014 |
| **S20HH14** | 2010 | Pakistan | Asia | ERR051783 | 6714_6_16 | | Shah et al. EID 2014 |
| **S16HH1** | 2010 | Pakistan | Asia | ERR051779 | 6714_6_12 | | Shah et al. EID 2014 |
| **S8KCH18** | 2010 | Pakistan | Asia | ERR051771 | 6714_6_4 | | Shah et al. EID 2014 |
| **F17KTH4** | 2010 | Pakistan | Asia | ERR051761 | 6714_5_17 | | Shah et al. EID 2014 |
| **F18KTH3** | 2010 | Pakistan | Asia | ERR051762 | 6714_5_18 | | Shah et al. EID 2014 |
| **S17HH3** | 2010 | Pakistan | Asia | ERR051780 | 6714_6_13 | | Shah et al. EID 2014 |
| **F12D1** | 2010 | Pakistan | Asia | ERR051756 | 6714_5_12 | | Shah et al. EID 2014 |
| **F5D38** | 2010 | Pakistan | Asia | ERR051749 | 6714_5_5 | | Shah et al. EID 2014 |
| **S1KCH15** | 2010 | Pakistan | Asia | ERR051764 | 6714_5_20 | | Shah et al. EID 2014 |
| **S21HH15** | 2010 | Pakistan | Asia | ERR051784 | 6714_6_17 | | Shah et al. EID 2014 |
| **S25R22** | 2010 | Pakistan | Asia | ERR051788 | 6714_6_21 | | Shah et al. EID 2014 |
| **S10P57** | 2010 | Pakistan | Asia | ERR051773 | 6714_6_6 | | Shah et al. EID 2014 |
| **F19KTH2** | 2010 | Pakistan | Asia | ERR051763 | 6714_5_19 | | Shah et al. EID 2014 |
| **S12P76** | 2010 | Pakistan | Asia | ERR051775 | 6714_6_8 | | Shah et al. EID 2014 |
| **S13P83** | 2010 | Pakistan | Asia | ERR051776 | 6714_6_9 | | Shah et al. EID 2014 |
| **S18HH4** | 2010 | Pakistan | Asia | ERR051781 | 6714_6_14 | | Shah et al. EID 2014 |
| **S19HH5** | 2010 | Pakistan | Asia | ERR051782 | 6714_6_15 | | Shah et al. EID 2014 |
| **F15KTH7** | 2010 | Pakistan | Asia | ERR051759 | 6714_5_15 | | Shah et al. EID 2014 |
| **F16KTH6** | 2010 | Pakistan | Asia | ERR051760 | 6714_5_16 | | Shah et al. EID 2014 |
| **H22** | 2022 | Haiti | North America | | H22 | | This Study |
| **D1** | 2021 | Bangladesh | Asia | | D1 | | This Study |

| | | | | | | | |
|---|---|---|---|---|---|---|---|
| **D2** | 2022 | Bangladesh | Asia | | D2 | | This Study |
| **D3** | 2022 | Bangladesh | Asia | | D3 | | This Study |
| **D4** | 2022 | Bangladesh | Asia | | D4 | | This Study |
| **EM1626** | 2011 | Bangladesh | Asia | SRR491153 | SRR491153 | | Unpublished EBI-ENA |
| **CP1040** | 2004 | Zambia | Africa | SRR227307 | SRR227307 | | Unpublished EBI-ENA |
| **CNRVC110078** | 2011 | Bangladesh | Asia | ERR2265590 | CNRVC110078 | | Weill et al. Nature 2019 |
| **CNRVC150256** | 2015 | Democratic Republic of the Congo | Africa | ERR2265661 | CNRVC150256 | | Weill et al. Nature 2019 |
| **CNRVC150071** | 2015 | Democratic Republic of the Congo | Africa | ERR2265651 | CNRVC150071 | | Weill et al. Nature 2019 |
| **CNRVC150056** | 2015 | Democratic Republic of the Congo | Africa | ERR2265650 | CNRVC150056 | | Weill et al. Nature 2019 |
| **THSTI_56712** | 2017 | India | Asia | ERR2270662 | THSTI_56712 | | Weill et al. Nature 2019 |
| **THSTI_56695** | 2017 | India | Asia | ERR2270661 | THSTI_56695 | | Weill et al. Nature 2019 |
| **THSTI_56650** | 2017 | India | Asia | ERR2270660 | THSTI_56650 | | Weill et al. Nature 2019 |
| **THSTI_55199** | 2017 | India | Asia | ERR2270659 | THSTI_55199 | | Weill et al. Nature 2019 |
| **THSTI_52665** | 2016 | India | Asia | ERR2270658 | THSTI_52665 | | Weill et al. Nature 2019 |
| **THSTI_52629** | 2016 | India | Asia | ERR2270657 | THSTI_52629 | | Weill et al. Nature 2019 |
| **THSTI_52588** | 2016 | India | Asia | ERR2270656 | THSTI_52588 | | Weill et al. Nature 2019 |
| **THSTI_52586** | 2016 | India | Asia | ERR2270655 | THSTI_52586 | | Weill et al. Nature 2019 |
| **THSTI_46073** | 2015 | India | Asia | ERR2269953 | THSTI_46073 | | Weill et al. Nature 2019 |
| **THSTI_45983** | 2015 | India | Asia | ERR2269952 | THSTI_45983 | | Weill et al. Nature 2019 |
| **THSTI_45869** | 2015 | India | Asia | ERR2269951 | THSTI_45869 | | Weill et al. Nature 2019 |
| **THSTI_45544** | 2015 | India | Asia | ERR2269950 | THSTI_45544 | | Weill et al. Nature 2019 |
| **THSTI_41081** | 2014 | India | Asia | ERR2269949 | THSTI_41081 | | Weill et al. Nature 2019 |
| **THSTI_41055** | 2014 | India | Asia | ERR2269948 | THSTI_41055 | | Weill et al. Nature 2019 |
| **THSTI_41049** | 2014 | India | Asia | ERR2269947 | THSTI_41049 | | Weill et al. Nature 2019 |
| **THSTI_36268** | 2013 | India | Asia | ERR2269946 | THSTI_36268 | | Weill et al. Nature 2019 |
| **THSTI_36136** | 2013 | India | Asia | ERR2269945 | THSTI_36136 | | Weill et al. Nature 2019 |

| | | | | | | | |
|---|---|---|---|---|---|---|---|
| **THSTI_36133** | 2013 | India | Asia | ERR2269944 | THSTI_36133 | | Weill et al. Nature 2019 |
| **THSTI_36124** | 2013 | India | Asia | ERR2269930 | THSTI_36124 | | Weill et al. Nature 2019 |
| **THSTI_33102** | 2012 | India | Asia | ERR2269929 | THSTI_33102 | | Weill et al. Nature 2019 |
| **THSTI_32676** | 2012 | India | Asia | ERR2269928 | THSTI_32676 | | Weill et al. Nature 2019 |
| **THSTI_31698** | 2012 | India | Asia | ERR2269927 | THSTI_31698 | | Weill et al. Nature 2019 |
| **THSTI_27071** | 2011 | India | Asia | ERR2269926 | THSTI_27071 | | Weill et al. Nature 2019 |
| **THSTI_26907** | 2011 | India | Asia | ERR2269925 | THSTI_26907 | | Weill et al. Nature 2019 |
| **THSTI_26871** | 2011 | India | Asia | ERR2269924 | THSTI_26871 | | Weill et al. Nature 2019 |
| **THSTI_26866** | 2011 | India | Asia | ERR2269923 | THSTI_26866 | | Weill et al. Nature 2019 |
| **THSTI_23835** | 2010 | India | Asia | ERR2269922 | THSTI_23835 | | Weill et al. Nature 2019 |
| **THSTI_23485** | 2010 | India | Asia | ERR2269921 | THSTI_23485 | | Weill et al. Nature 2019 |
| **THSTI_20370** | 2010 | India | Asia | ERR2269808 | THSTI_20370 | | Weill et al. Nature 2019 |
| **CNRVC150140** | 2015 | India | Asia | ERR2265652 | CNRVC150140 | | Weill et al. Nature 2019 |
| **CNRVC140176** | 2014 | India | Asia | ERR2265649 | CNRVC140176 | | Weill et al. Nature 2019 |
| **CNRVC170252** | 2015 | Iran | Asia | ERR2269838 | CNRVC170252 | | Weill et al. Nature 2019 |
| **CNRVC170250** | 2013 | Iran | Asia | ERR2269837 | CNRVC170250 | | Weill et al. Nature 2019 |
| **CNRVC170249** | 2013 | Iran | Asia | ERR2269836 | CNRVC170249 | | Weill et al. Nature 2019 |
| **CNRVC170248** | 2012 | Iran | Asia | ERR2269835 | CNRVC170248 | | Weill et al. Nature 2019 |
| **CNRVC160018** | 2007 | Iraq | Asia | ERR2265666 | CNRVC160018_TCATTC_L001 | | Weill et al. Nature 2019 |
| **CNRVC160017** | 2007 | Iraq | Asia | ERR2265665 | CNRVC160017_TATAAT_L001 | | Weill et al. Nature 2019 |
| **CNRVC160016** | 2007 | Iraq | Asia | ERR2265664 | CNRVC160016_TACAGC_L001 | | Weill et al. Nature 2019 |
| **CNRVC160014** | 2007 | Iraq | Asia | ERR2265663 | CNRVC160014_GCGCTA_L001 | | Weill et al. Nature 2019 |
| **CNRVC160013** | 2007 | Iraq | Asia | ERR2265662 | CNRVC160013_CTCAGA_L001 | | Weill et al. Nature 2019 |
| **CNRVC150181** | 2015 | Iraq | Asia | ERR2265660 | CNRVC150181_CCAACA_L001 | | Weill et al. Nature 2019 |
| **CNRVC150177** | 2015 | Iraq | Asia | ERR2265659 | CNRVC150177_CACTCA_L001 | | Weill et al. Nature 2019 |
| **CNRVC150171** | 2015 | Iraq | Asia | ERR2265658 | CNRVC150171_ATGAGC_L001 | | Weill et al. Nature 2019 |

| | | | | | | | |
|---|---|---|---|---|---|---|---|
| **CNRVC150170** | 2015 | Iraq | Asia | ERR2265657 | CNRVC150170_ACTGAT_L001 | | Weill et al. Nature 2019 |
| **CNRVC150169** | 2015 | Iraq | Asia | ERR2265656 | CNRVC150169_GGTAGC_L001 | | Weill et al. Nature 2019 |
| **CNRVC150168** | 2015 | Iraq | Asia | ERR2265655 | CNRVC150168_GAGTGG_L001 | | Weill et al. Nature 2019 |
| **31_1** | 2015 | Kenya | Africa | ERR2265589 | VC_31_1 | | Weill et al. Nature 2019 |
| **4621STDY6714780** | 2016 | Kenya | Africa | ERS1572815 | 22204_7_320 | | Weill et al. Nature 2019 |
| **4621STDY6714778** | 2010 | Kenya | Africa | ERS1572813 | 22204_7_318 | | Weill et al. Nature 2019 |
| **4621STDY6714774** | 2012 | Kenya | Africa | ERS1572809 | 22204_7_314 | | Weill et al. Nature 2019 |
| **4621STDY6714768** | 2012 | Kenya | Africa | ERS1572803 | 22204_7_308 | | Weill et al. Nature 2019 |
| **4621STDY6714763** | 2012 | Kenya | Africa | ERS1572798 | 22204_7_303 | | Weill et al. Nature 2019 |
| **4621STDY6714758** | 2015 | Kenya | Africa | ERS1572793 | 22204_7_298 | | Weill et al. Nature 2019 |
| **4621STDY6714750** | 2015 | Kenya | Africa | ERS1572785 | 22204_7_290 | | Weill et al. Nature 2019 |
| **4621STDY6714749** | 2015 | Kenya | Africa | ERS1572784 | 22204_7_289 | | Weill et al. Nature 2019 |
| **4621STDY6714748** | 2015 | Kenya | Africa | ERS1572783 | 22204_7_288 | | Weill et al. Nature 2019 |
| **CNRVC170165** | 2017 | South Sudan | Africa | ERR2265673 | CNRVC170165 | | Weill et al. Nature 2019 |
| **CNRVC170164** | 2017 | South Sudan | Africa | ERR2265672 | CNRVC170164 | | Weill et al. Nature 2019 |
| **CNRVC170161** | 2017 | South Sudan | Africa | ERR2265671 | CNRVC170161 | | Weill et al. Nature 2019 |
| **CNRVC170160** | 2017 | South Sudan | Africa | ERR2265670 | CNRVC170160 | | Weill et al. Nature 2019 |
| **CNRVC170159** | 2017 | South Sudan | Africa | ERR2265669 | CNRVC170159 | | Weill et al. Nature 2019 |
| CNRVC160462 | 2016 | South Sudan | Africa | ERR2265668 | CNRVC160462 | | Weill et al. Nature 2019 |
| **CNRVC160461** | 2016 | South Sudan | Africa | ERR2265667 | CNRVC160461 | | Weill et al. Nature 2019 |
| **CNRVC150165** | 2015 | South Sudan | Africa | ERR2265654 | CNRVC150165 | | Weill et al. Nature 2019 |
| **CNRVC150141** | 2015 | South Sudan | Africa | ERR2265653 | CNRVC150141 | | Weill et al. Nature 2019 |
| **CNRVC140079** | 2014 | South Sudan | Africa | ERR2265648 | CNRVC140079 | | Weill et al. Nature 2019 |
| **CNRVC140077** | 2014 | South Sudan | Africa | ERR2265647 | CNRVC140077 | | Weill et al. Nature 2019 |

| | | | | | | | |
|---|---|---|---|---|---|---|---|
| **CNRVC140072** | 2014 | South Sudan | Africa | ERR2265646 | CNRVC140072 | | Weill et al. Nature 2019 |
| **CNRVC140069** | 2014 | South Sudan | Africa | ERR2265645 | CNRVC140069 | | Weill et al. Nature 2019 |
| **CNRVC140054** | 2014 | South Sudan | Africa | ERR2265591 | CNRVC140054 | | Weill et al. Nature 2019 |
| **CNRVC170242** | 2017 | Yemen | Asia | ERR2269834 | CNRVC170242 | | Weill et al. Nature 2019 |
| **CNRVC170241** | 2017 | Yemen | Asia | ERR2269833 | CNRVC170241 | | Weill et al. Nature 2019 |
| **CNRVC170240** | 2017 | Yemen | Asia | ERR2269832 | CNRVC170240 | | Weill et al. Nature 2019 |
| **CNRVC170208** | 2017 | Yemen | Asia | ERR2269811 | CNRVC170208 | | Weill et al. Nature 2019 |
| **CNRVC170207** | 2017 | Yemen | Asia | ERR2269810 | CNRVC170207 | | Weill et al. Nature 2019 |
| **CNRVC170206** | 2017 | Yemen | Asia | ERR2269809 | CNRVC170206 | | Weill et al. Nature 2019 |
| **CNRVC170205** | 2017 | Yemen | Asia | ERR2269718 | CNRVC170205 | | Weill et al. Nature 2019 |
| **CNRVC170204** | 2017 | Yemen | Asia | ERR2269717 | CNRVC170204 | | Weill et al. Nature 2019 |
| **CNRVC170203** | 2017 | Yemen | Asia | ERR2269716 | CNRVC170203 | | Weill et al. Nature 2019 |
| **CNRVC170202** | 2017 | Yemen | Asia | ERR2269715 | CNRVC170202 | | Weill et al. Nature 2019 |
| **CNRVC170201** | 2017 | Yemen | Asia | ERR2269714 | CNRVC170201 | | Weill et al. Nature 2019 |
| **CNRVC170200** | 2017 | Yemen | Asia | ERR2269713 | CNRVC170200 | | Weill et al. Nature 2019 |
| **CNRVC170199** | 2017 | Yemen | Asia | ERR2269712 | CNRVC170199 | | Weill et al. Nature 2019 |
| **CNRVC170198** | 2017 | Yemen | Asia | ERR2269711 | CNRVC170198 | | Weill et al. Nature 2019 |
| **CNRVC170197** | 2017 | Yemen | Asia | ERR2269710 | CNRVC170197 | | Weill et al. Nature 2019 |
| **CNRVC170196** | 2017 | Yemen | Asia | ERR2269709 | CNRVC170196 | | Weill et al. Nature 2019 |
| **CNRVC170195** | 2017 | Yemen | Asia | ERR2269650 | CNRVC170195 | | Weill et al. Nature 2019 |
| **CNRVC170194** | 2017 | Yemen | Asia | ERR2269649 | CNRVC170194 | | Weill et al. Nature 2019 |
| **CNRVC170193** | 2017 | Yemen | Asia | ERR2269648 | CNRVC170193 | | Weill et al. Nature 2019 |
| **CNRVC170192** | 2017 | Yemen | Asia | ERR2269647 | CNRVC170192 | | Weill et al. Nature 2019 |
| **CNRVC170191** | 2017 | Yemen | Asia | ERR2269646 | CNRVC170191 | | Weill et al. Nature 2019 |
| **CNRVC170190** | 2017 | Yemen | Asia | ERR2269645 | CNRVC170190 | | Weill et al. Nature 2019 |

| | | | | | | | |
|---|---|---|---|---|---|---|---|
| **CNRVC170189** | 2017 | Yemen | Asia | ERR2269644 | CNRVC170189 | | Weill et al. Nature 2019 |
| **CNRVC170188** | 2017 | Yemen | Asia | ERR2269643 | CNRVC170188 | | Weill et al. Nature 2019 |
| **CNRVC170187** | 2017 | Yemen | Asia | ERR2269642 | CNRVC170187 | | Weill et al. Nature 2019 |
| **CNRVC170186** | 2017 | Yemen | Asia | ERR2269641 | CNRVC170186 | | Weill et al. Nature 2019 |
| **CNRVC170185** | 2017 | Yemen | Asia | ERR2269640 | CNRVC170185 | | Weill et al. Nature 2019 |
| **CNRVC170184** | 2017 | Yemen | Asia | ERR2269622 | CNRVC170184 | | Weill et al. Nature 2019 |
| **CNRVC170183** | 2017 | Yemen | Asia | ERR2269621 | CNRVC170183 | | Weill et al. Nature 2019 |
| **CNRVC170182** | 2017 | Yemen | Asia | ERR2269620 | CNRVC170182 | | Weill et al. Nature 2019 |
| **CNRVC170181** | 2017 | Yemen | Asia | ERR2269619 | CNRVC170181 | | Weill et al. Nature 2019 |
| **CNRVC170180** | 2017 | Yemen | Asia | ERR2269618 | CNRVC170180 | | Weill et al. Nature 2019 |
| **CNRVC170179** | 2017 | Yemen | Asia | ERR2269617 | CNRVC170179 | | Weill et al. Nature 2019 |
| **CNRVC170178** | 2017 | Yemen | Asia | ERR2269616 | CNRVC170178 | | Weill et al. Nature 2019 |
| **CNRVC170177** | 2016 | Yemen | Asia | ERR2269615 | CNRVC170177 | | Weill et al. Nature 2019 |
| **CNRVC170176** | 2016 | Yemen | Asia | ERR2269614 | CNRVC170176 | | Weill et al. Nature 2019 |
| **CNRVC170175** | 2016 | Yemen | Asia | ERR2269613 | CNRVC170175 | | Weill et al. Nature 2019 |
| **CNRVC170174** | 2016 | Yemen | Asia | ERR2265678 | CNRVC170174 | | Weill et al. Nature 2019 |
| **CNRVC170173** | 2016 | Yemen | Asia | ERR2265677 | CNRVC170173 | | Weill et al. Nature 2019 |
| **CNRVC170170** | 2016 | Yemen | Asia | ERR2265676 | CNRVC170170 | | Weill et al. Nature 2019 |
| **CNRVC170169** | 2016 | Yemen | Asia | ERR2265675 | CNRVC170169 | | Weill et al. Nature 2019 |
| **CNRVC170168** | 2016 | Yemen | Asia | ERR2265674 | CNRVC170168 | | Weill et al. Nature 2019 |
| **CNRVC990299** | 1999 | Afghanistan | Asia | ERR1879649 | CNRVC990299_CCGTCC_L001 | | Weill et al. Science 2017 |
| **CNRVC990298** | 1999 | Afghanistan | Asia | ERR1879648 | CNRVC990298_ATGTCA_L001 | | Weill et al. Science 2017 |
| **CNRVC940149** | 1994 | Albania | Europe | ERR1879577 | CNRVC940149_AGTCAA_L002 | | Weill et al. Science 2017 |
| **CNRVC070154** | 1994 | Algeria | Africa | ERR1878560 | CNRVC070154_GGCTAC_L001 | | Weill et al. Science 2017 |
| **CNRVC070120** | 1994 | Algeria | Africa | ERR1878555 | CNRVC070120_TAGCTT_L001 | | Weill et al. Science 2017 |

| | | | | | | | |
|---|---|---|---|---|---|---|---|
| **CNRVC970008** | 1990 | Algeria | Africa | ERR976476 | 16244_7_25 | | Weill et al. Science 2017 |
| **CNRVC950418** | 1974 | Algeria | Africa | ERR976444 | 16244_6_87 | | Weill et al. Science 2017 |
| **CNRVC930187** | 1982 | Algeria | Africa | ERR976413 | 16244_6_56 | | Weill et al. Science 2017 |
| **CNRVC930172** | 1986 | Algeria | Africa | ERR976412 | 16244_6_55 | | Weill et al. Science 2017 |
| **CNRVC930169** | 1986 | Algeria | Africa | ERR976409 | 16244_6_52 | | Weill et al. Science 2017 |
| **CNRVC930168** | 1982 | Algeria | Africa | ERR976408 | 16244_6_51 | | Weill et al. Science 2017 |
| **CNRVC930167** | 1974 | Algeria | Africa | ERR976407 | 16244_6_50 | | Weill et al. Science 2017 |
| **CNRVC930166** | 1975 | Algeria | Africa | ERR976406 | 16244_6_49 | | Weill et al. Science 2017 |
| **CNRVC930164** | 1983 | Algeria | Africa | ERR976405 | 16244_6_48 | | Weill et al. Science 2017 |
| **CNRVC920172** | 1987 | Algeria | Africa | ERR976395 | 16244_6_38 | | Weill et al. Science 2017 |
| **RKI-ZBS2-CH19** | 1972 | Angola | Africa | ERR1880793 | RKI-ZBS2-CH19_GCGCTA_L002 | | Weill et al. Science 2017 |
| **CNRVC950002** | 1995 | Angola | Africa | ERR998664 | 16356_8_9 | | Weill et al. Science 2017 |
| **CNRVC950007** | 1995 | Angola | Africa | ERR998665 | 16356_8_10 | | Weill et al. Science 2017 |
| **CNRVC940012** | 1994 | Angola | Africa | ERR976533 | 16244_7_82 | | Weill et al. Science 2017 |
| **CNRVC940011** | 1994 | Angola | Africa | ERR976532 | 16244_7_81 | | Weill et al. Science 2017 |
| **CNRVC920027** | 1992 | Angola | Africa | ERR976522 | 16244_7_71 | | Weill et al. Science 2017 |
| **CNRVC950569** | 1970 | Angola | Africa | ERR976457 | 16244_7_6 | | Weill et al. Science 2017 |
| **CNRVC140132** | 1988 | Angola | Africa | ERR976503 | 16244_7_52 | | Weill et al. Science 2017 |
| **CNRVC140131** | 1988 | Angola | Africa | ERR976502 | 16244_7_51 | | Weill et al. Science 2017 |
| **CNRVC140130** | 1988 | Angola | Africa | ERR976501 | 16244_7_50 | | Weill et al. Science 2017 |
| **CNRVC950568** | 1970 | Angola | Africa | ERR976456 | 16244_7_5 | | Weill et al. Science 2017 |
| **CNRVC140129** | 1988 | Angola | Africa | ERR976500 | 16244_7_49 | | Weill et al. Science 2017 |
| **CNRVC140128** | 1988 | Angola | Africa | ERR976499 | 16244_7_48 | | Weill et al. Science 2017 |
| **CNRVC140127** | 1988 | Angola | Africa | ERR976498 | 16244_7_47 | | Weill et al. Science 2017 |
| **CNRVC920194** | 1990 | Angola | Africa | ERR976396 | 16244_6_39 | | Weill et al. Science 2017 |

| | | | | | | | |
|---|---|---|---|---|---|---|---|
| **CNRVC900105** | 1990 | Angola | Africa | ERR976391 | 16244_6_34 | | Weill et al. Science 2017 |
| **CNRVC100183** | 2010 | Benin | Africa | ERR1878587 | CNRVC100183 | | Weill et al. Science 2017 |
| **CNRVC100176** | 2010 | Benin | Africa | ERR1878586 | CNRVC100176 | | Weill et al. Science 2017 |
| **CNRVC070164** | 2007 | Benin | Africa | ERR1878561 | CNRVC070164 | | Weill et al. Science 2017 |
| **CNRVC050140** | 2005 | Benin | Africa | ERR1878134 | CNRVC050140 | | Weill et al. Science 2017 |
| **CNRVC050139** | 2005 | Benin | Africa | ERR1878133 | CNRVC050139 | | Weill et al. Science 2017 |
| **CNRVC040132** | 2004 | Benin | Africa | ERR1878110 | CNRVC040132 | | Weill et al. Science 2017 |
| **CNRVC040127** | 2004 | Benin | Africa | ERR1878109 | CNRVC040127 | | Weill et al. Science 2017 |
| **CNRVC030084** | 2003 | Benin | Africa | ERR1878092 | CNRVC030084 | | Weill et al. Science 2017 |
| **CNRVC030082** | 2003 | Benin | Africa | ERR1878091 | CNRVC030082 | | Weill et al. Science 2017 |
| **CNRVC020367** | 2002 | Benin | Africa | ERR1877957 | CNRVC020367 | | Weill et al. Science 2017 |
| **CNRVC020361** | 2002 | Benin | Africa | ERR1877956 | CNRVC020361 | | Weill et al. Science 2017 |
| **CNRVC970152** | 1997 | Benin | Africa | ERR998750 | 16356_8_95 | | Weill et al. Science 2017 |
| **CNRVC970147** | 1997 | Benin | Africa | ERR998749 | 16356_8_94 | | Weill et al. Science 2017 |
| **CNRVC960579** | 1970 | Benin | Africa | ERR976460 | 16244_7_9 | | Weill et al. Science 2017 |
| **CNRVC960575** | 1970 | Benin | Africa | ERR976459 | 16244_7_8 | | Weill et al. Science 2017 |
| **CNRVC910267** | 1991 | Benin | Africa | ERR976515 | 16244_7_64 | | Weill et al. Science 2017 |
| **CNRVC910266** | 1991 | Benin | Africa | ERR976514 | 16244_7_63 | | Weill et al. Science 2017 |
| **CNRVC910036** | 1991 | Benin | Africa | ERR976513 | 16244_7_62 | | Weill et al. Science 2017 |
| **CNRVC980046** | 1985 | Benin | Africa | ERR976483 | 16244_7_32 | | Weill et al. Science 2017 |
| **CNRVC990234** | 1999 | Burkina Faso | Africa | ERR1879646 | CNRVC990234_CTATAC_L002 | | Weill et al. Science 2017 |
| **CNRVC070334** | 2005 | Burkina Faso | Africa | ERR1878563 | CNRVC070334 | | Weill et al. Science 2017 |
| **CNRVC070316** | 2005 | Burkina Faso | Africa | ERR1878562 | CNRVC070316 | | Weill et al. Science 2017 |
| **CNRVC010206** | 2001 | Burkina Faso | Africa | ERR1877949 | CNRVC010206 | | Weill et al. Science 2017 |
| **CNRVC010205** | 2001 | Burkina Faso | Africa | ERR1877948 | CNRVC010205 | | Weill et al. Science 2017 |

| | | | | | | | |
|---|---|---|---|---|---|---|---|
| **CNRVC950806** | 1995 | Burkina Faso | Africa | ERR998688 | 16356_8_33 | | Weill et al. Science 2017 |
| **CNRVC950801** | 1995 | Burkina Faso | Africa | ERR998687 | 16356_8_32 | | Weill et al. Science 2017 |
| **CNRVC950702** | 1995 | Burkina Faso | Africa | ERR998672 | 16356_8_17 | | Weill et al. Science 2017 |
| **CNRVC950700** | 1995 | Burkina Faso | Africa | ERR998671 | 16356_8_16 | | Weill et al. Science 2017 |
| **CNRVC980360** | 1998 | Burkina Faso | Africa | ERR976573 | 16244_8_27 | | Weill et al. Science 2017 |
| **CNRVC980359** | 1998 | Burkina Faso | Africa | ERR976572 | 16244_8_26 | | Weill et al. Science 2017 |
| **CNRVC980049** | 1984 | Burkina Faso | Africa | ERR976486 | 16244_7_35 | | Weill et al. Science 2017 |
| **CNRVC980048** | 1984 | Burkina Faso | Africa | ERR976485 | 16244_7_34 | | Weill et al. Science 2017 |
| **CNRVC980040** | 1984 | Burkina Faso | Africa | ERR976479 | 16244_7_28 | | Weill et al. Science 2017 |
| **CNRVC950405** | 1974 | Burkina Faso | Africa | ERR976443 | 16244_6_86 | | Weill et al. Science 2017 |
| **CNRVC950396** | 1974 | Burkina Faso | Africa | ERR976442 | 16244_6_85 | | Weill et al. Science 2017 |
| **CNRVC950106** | 1974 | Burkina Faso | Africa | ERR976419 | 16244_6_62 | | Weill et al. Science 2017 |
| **CNRVC950105** | 1974 | Burkina Faso | Africa | ERR976418 | 16244_6_61 | | Weill et al. Science 2017 |
| **CNRVC970056** | 1997 | Burundi | Africa | ERR1879636 | CNRVC970056_CATTTT_L002 | | Weill et al. Science 2017 |
| **CNRVC010062** | 2001 | Burundi | Africa | ERR1877942 | CNRVC010062 | | Weill et al. Science 2017 |
| **CNRVC010061** | 2001 | Burundi | Africa | ERR1877941 | CNRVC010061 | | Weill et al. Science 2017 |
| **C8466** | 1992 | Burundi | Africa | ERR1877618 | CDCC8466_rep | | Weill et al. Science 2017 |
| **C8465** | 1992 | Burundi | Africa | ERR1877617 | CDCC8465_rep | | Weill et al. Science 2017 |
| **CNRVC930425** | 1993 | Burundi | Africa | ERR976530 | 16244_7_79 | | Weill et al. Science 2017 |
| **CNRVC930417** | 1993 | Burundi | Africa | ERR976529 | 16244_7_78 | | Weill et al. Science 2017 |
| **CNRVC990194** | 1999 | Cambodia | Asia | ERR1879644 | CNRVC990194_CGGAAT_L002 | | Weill et al. Science 2017 |
| **CNRVC930067** | 1963 | Cambodia | Asia | ERR1879567 | CNRVC930067_GCCAAT_L002 | | Weill et al. Science 2017 |
| **CNRVC910040** | 1991 | Cambodia | Asia | ERR1879548 | CNRVC910040_CAGATC_L001 | | Weill et al. Science 2017 |
| **CNRVC150239** | 1993 | Cambodia | Asia | ERR1879538 | CNRVC150239_CGGAAT_L001 | | Weill et al. Science 2017 |
| **CNRVC990329** | 1999 | Cameroon | Africa | ERR1879650 | CNRVC990329_CGATGT_L001 | | Weill et al. Science 2017 |

| | | | | | | | |
|---|---|---|---|---|---|---|---|
| **CNRVC110128** | 2010 | Cameroon | Africa | ERR1878598 | CNRVC110128 | | Weill et al. Science 2017 |
| **CNRVC110120** | 2011 | Cameroon | Africa | ERR1878597 | CNRVC110120 | | Weill et al. Science 2017 |
| **CNRVC110113** | 2011 | Cameroon | Africa | ERR1878596 | CNRVC110113 | | Weill et al. Science 2017 |
| **CNRVC110096** | 2010 | Cameroon | Africa | ERR1878595 | CNRVC110096 | | Weill et al. Science 2017 |
| **CNRVC100186** | 2010 | Cameroon | Africa | ERR1878588 | CNRVC100186 | | Weill et al. Science 2017 |
| **CNRVC090182** | 2009 | Cameroon | Africa | ERR1878580 | CNRVC090182 | | Weill et al. Science 2017 |
| **CNRVC080764** | 2008 | Cameroon | Africa | ERR1878574 | CNRVC080764 | | Weill et al. Science 2017 |
| **CNRVC080762** | 2008 | Cameroon | Africa | ERR1878573 | CNRVC080762 | | Weill et al. Science 2017 |
| **CNRVC060111** | 2006 | Cameroon | Africa | ERR1878152 | CNRVC060111 | | Weill et al. Science 2017 |
| **CNRVC050011** | 2005 | Cameroon | Africa | ERR1878129 | CNRVC050011 | | Weill et al. Science 2017 |
| **CNRVC050008** | 2005 | Cameroon | Africa | ERR1878128 | CNRVC050008 | | Weill et al. Science 2017 |
| **CNRVC040110** | 2004 | Cameroon | Africa | ERR1878108 | CNRVC040110 | | Weill et al. Science 2017 |
| **CNRVC040082** | 2002 | Cameroon | Africa | ERR1878107 | CNRVC040082 | | Weill et al. Science 2017 |
| **CNRVC040074** | 2001 | Cameroon | Africa | ERR1878106 | CNRVC040074 | | Weill et al. Science 2017 |
| **CNRVC040061** | 2004 | Cameroon | Africa | ERR1878105 | CNRVC040061 | | Weill et al. Science 2017 |
| **CNRVC010221** | 2001 | Cameroon | Africa | ERR1877950 | CNRVC010221 | | Weill et al. Science 2017 |
| **CNRVC010023** | 2000 | Cameroon | Africa | ERR1877645 | CNRVC010023 | | Weill et al. Science 2017 |
| **CNRVC000323** | 2000 | Cameroon | Africa | ERR1877643 | CNRVC000323 | | Weill et al. Science 2017 |
| **F135** | 1993 | Cameroon | Africa | ERR1879651 | CDCF135_rep | | Weill et al. Science 2017 |
| **CNRVC970036** | 1997 | Cameroon | Africa | ERR998731 | 16356_8_76 | | Weill et al. Science 2017 |
| **CNRVC970024** | 1997 | Cameroon | Africa | ERR998728 | 16356_8_73 | | Weill et al. Science 2017 |
| **CNRVC970022** | 1997 | Cameroon | Africa | ERR998727 | 16356_8_72 | | Weill et al. Science 2017 |
| **CNRVC970019** | 1997 | Cameroon | Africa | ERR998726 | 16356_8_71 | | Weill et al. Science 2017 |
| **CNRVC970014** | 1997 | Cameroon | Africa | ERR998725 | 16356_8_70 | | Weill et al. Science 2017 |
| **CNRVC960293** | 1996 | Cameroon | Africa | ERR998712 | 16356_8_57 | | Weill et al. Science 2017 |

| | | | | | | | |
|---|---|---|---|---|---|---|---|
| **CNRVC960246** | 1996 | Cameroon | Africa | ERR998703 | 16356_8_48 | | Weill et al. Science 2017 |
| **CNRVC950011** | 1995 | Cameroon | Africa | ERR998666 | 16356_8_11 | | Weill et al. Science 2017 |
| **CNRVC980022** | 1997 | Cameroon | Africa | ERR976555 | 16244_8_9 | | Weill et al. Science 2017 |
| **CNRVC980021** | 1997 | Cameroon | Africa | ERR976554 | 16244_8_8 | | Weill et al. Science 2017 |
| **CNRVC980398** | 1998 | Cameroon | Africa | ERR976579 | 16244_8_33 | | Weill et al. Science 2017 |
| **CNRVC980397** | 1998 | Cameroon | Africa | ERR976578 | 16244_8_32 | | Weill et al. Science 2017 |
| **CNRVC980396** | 1998 | Cameroon | Africa | ERR976577 | 16244_8_31 | | Weill et al. Science 2017 |
| **CNRVC980395** | 1998 | Cameroon | Africa | ERR976576 | 16244_8_30 | | Weill et al. Science 2017 |
| **CNRVC980328** | 1998 | Cameroon | Africa | ERR976570 | 16244_8_24 | | Weill et al. Science 2017 |
| **CNRVC980323** | 1998 | Cameroon | Africa | ERR976568 | 16244_8_22 | | Weill et al. Science 2017 |
| **CNRVC980061** | 1998 | Cameroon | Africa | ERR976562 | 16244_8_16 | | Weill et al. Science 2017 |
| **CNRVC980060** | 1998 | Cameroon | Africa | ERR976561 | 16244_8_15 | | Weill et al. Science 2017 |
| **CNRVC950143** | 1970 | Cameroon | Africa | ERR976424 | 16244_6_67 | | Weill et al. Science 2017 |
| **CNRVC950142** | 1970 | Cameroon | Africa | ERR976423 | 16244_6_66 | | Weill et al. Science 2017 |
| **CNRVC950141** | 1970 | Cameroon | Africa | ERR976422 | 16244_6_65 | | Weill et al. Science 2017 |
| **CNRVC950140** | 1970 | Cameroon | Africa | ERR976421 | 16244_6_64 | | Weill et al. Science 2017 |
| **CNRVC950138** | 1970 | Cameroon | Africa | ERR976420 | 16244_6_63 | | Weill et al. Science 2017 |
| **CNRVC930042** | 1970 | Cameroon | Africa | ERR976400 | 16244_6_43 | | Weill et al. Science 2017 |
| **CNRVC110272** | 2011 | Central African Republic | Africa | ERR1878603 | CNRVC110272 | | Weill et al. Science 2017 |
| **CNRVC110266** | 2011 | Central African Republic | Africa | ERR1878602 | CNRVC110266 | | Weill et al. Science 2017 |
| **CNRVC970143** | 1997 | Central African Republic | Africa | ERR998748 | 16356_8_93 | | Weill et al. Science 2017 |
| **CNRVC970141** | 1997 | Central African Republic | Africa | ERR998747 | 16356_8_92 | | Weill et al. Science 2017 |
| **CNRVC970127** | 1997 | Central African Republic | Africa | ERR998745 | 16356_8_90 | | Weill et al. Science 2017 |
| **CNRVC970126** | 1997 | Central African Republic | Africa | ERR998744 | 16356_8_89 | | Weill et al. Science 2017 |
| **CNRVC970125** | 1997 | Central African Republic | Africa | ERR998743 | 16356_8_88 | | Weill et al. Science 2017 |

| | | | | | | | |
|---|---|---|---|---|---|---|---|
| **CNRVC970123** | 1997 | Central African Republic | Africa | ERR998742 | 16356_8_87 | | Weill et al. Science 2017 |
| **CNRVC970112** | 1997 | Central African Republic | Africa | ERR998740 | 16356_8_85 | | Weill et al. Science 2017 |
| **CNRVC970110** | 1997 | Central African Republic | Africa | ERR998739 | 16356_8_84 | | Weill et al. Science 2017 |
| **CNRVC970109** | 1997 | Central African Republic | Africa | ERR998738 | 16356_8_83 | | Weill et al. Science 2017 |
| **CNRVC970108** | 1997 | Central African Republic | Africa | ERR998737 | 16356_8_82 | | Weill et al. Science 2017 |
| **CNRVC970079** | 1997 | Central African Republic | Africa | ERR998736 | 16356_8_81 | | Weill et al. Science 2017 |
| **CNRVC970075** | 1997 | Central African Republic | Africa | ERR998735 | 16356_8_80 | | Weill et al. Science 2017 |
| **CNRVC980374** | 1998 | Chad | Africa | ERR1879638 | CNRVC980374_TCCCGA_L002 | | Weill et al. Science 2017 |
| **CNRVC110241** | 2011 | Chad | Africa | ERR1878600 | CNRVC110241 | | Weill et al. Science 2017 |
| **CNRVC110230** | 2011 | Chad | Africa | ERR1878599 | CNRVC110230 | | Weill et al. Science 2017 |
| **CNRVC100246** | 2010 | Chad | Africa | ERR1878591 | CNRVC100246 | | Weill et al. Science 2017 |
| **CNRVC100240** | 2010 | Chad | Africa | ERR1878590 | CNRVC100240 | | Weill et al. Science 2017 |
| **CNRVC010203** | 2001 | Chad | Africa | ERR1877947 | CNRVC010203 | | Weill et al. Science 2017 |
| **CNRVC010123** | 2001 | Chad | Africa | ERR1877946 | CNRVC010123_ACTTGA_L001 | | Weill et al. Science 2017 |
| **CNRVC010120** | 2001 | Chad | Africa | ERR1877945 | CNRVC010120 | | Weill et al. Science 2017 |
| **CNRVC940184** | 1994 | Chad | Africa | ERR998662 | 16356_8_7 | | Weill et al. Science 2017 |
| **CNRVC960273** | 1996 | Chad | Africa | ERR998711 | 16356_8_56 | | Weill et al. Science 2017 |
| **CNRVC960265** | 1996 | Chad | Africa | ERR998708 | 16356_8_53 | | Weill et al. Science 2017 |
| **CNRVC960243** | 1996 | Chad | Africa | ERR998702 | 16356_8_47 | | Weill et al. Science 2017 |
| **CNRVC960234** | 1996 | Chad | Africa | ERR998701 | 16356_8_46 | | Weill et al. Science 2017 |
| **CNRVC960228** | 1996 | Chad | Africa | ERR998700 | 16356_8_45 | | Weill et al. Science 2017 |
| **CNRVC940163** | 1994 | Chad | Africa | ERR998656 | 16356_8_1 | | Weill et al. Science 2017 |
| **CNRVC950366** | 1974 | Chad | Africa | ERR976440 | 16244_6_83 | | Weill et al. Science 2017 |
| **CNRVC950364** | 1972 | Chad | Africa | ERR976439 | 16244_6_82 | | Weill et al. Science 2017 |
| **CNRVC950360** | 1974 | Chad | Africa | ERR976438 | 16244_6_81 | | Weill et al. Science 2017 |

| | | | | | | | |
|---|---|---|---|---|---|---|---|
| **CNRVC950358** | 1974 | Chad | Africa | ERR976437 | 16244_6_80 | | Weill et al. Science 2017 |
| **CNRVC950087** | 1972 | Chad | Africa | ERR976416 | 16244_6_59 | | Weill et al. Science 2017 |
| **CNRVC950040** | 1972 | Chad | Africa | ERR976415 | 16244_6_58 | | Weill et al. Science 2017 |
| **CNRVC930046** | 1971 | Chad | Africa | ERR976401 | 16244_6_44 | | Weill et al. Science 2017 |
| **CNRVC930061** | 1961 | China | Asia | ERR1879565 | CNRVC930061_TGACCA_L002 | | Weill et al. Science 2017 |
| **CNRVC030594** | 2003 | Comoros | Africa | ERR1878104 | CNRVC030594 | | Weill et al. Science 2017 |
| **CNRVC030593** | 2003 | Comoros | Africa | ERR1878103 | CNRVC030593 | | Weill et al. Science 2017 |
| **CNRVC020003** | 2002 | Comoros | Africa | ERR1877953 | CNRVC020003 | | Weill et al. Science 2017 |
| **CNRVC010045** | 2001 | Comoros | Africa | ERR1877648 | CNRVC010045 | | Weill et al. Science 2017 |
| **CNRVC010008** | 2001 | Comoros | Africa | ERR1877644 | CNRVC010008_GCCAAT_L001 | | Weill et al. Science 2017 |
| **CNRVC000085** | 2000 | Comoros | Africa | ERR1877640 | CNRVC000085 | | Weill et al. Science 2017 |
| **CNRVC980029** | 1998 | Comoros | Africa | ERR976558 | 16244_8_12 | | Weill et al. Science 2017 |
| **CNRVC980026** | 1998 | Comoros | Africa | ERR976557 | 16244_8_11 | | Weill et al. Science 2017 |
| **CNRVC980023** | 1998 | Comoros | Africa | ERR976556 | 16244_8_10 | | Weill et al. Science 2017 |
| **CNRVC950543** | 1975 | Comoros | Africa | ERR976453 | 16244_7_2 | | Weill et al. Science 2017 |
| **CNRVC950540** | 1975 | Comoros | Africa | ERR976452 | 16244_7_1 | | Weill et al. Science 2017 |
| **CNRVC950535** | 1975 | Comoros | Africa | ERR976451 | 16244_6_94 | | Weill et al. Science 2017 |
| **CNRVC150245** | 1988 | Cote d'Ivoire | Africa | ERR1879542 | CNRVC150245_GCGCTA_L001 | | Weill et al. Science 2017 |
| **CNRVC150231** | 1988 | Cote d'Ivoire | Africa | ERR1879535 | CNRVC150231_CATGGC_L001 | | Weill et al. Science 2017 |
| **CNRVC150227** | 1988 | Cote d'Ivoire | Africa | ERR1879435 | CNRVC150227_CACCGG_L001 | | Weill et al. Science 2017 |
| **CNRVC150210** | 1988 | Cote d'Ivoire | Africa | ERR1879384 | CNRVC150210_AGTCAA_L001 | | Weill et al. Science 2017 |
| **CNRVC080497** | 2006 | Cote d'Ivoire | Africa | ERR1878572 | CNRVC080497 | | Weill et al. Science 2017 |
| **CNRVC080496** | 2006 | Cote d'Ivoire | Africa | ERR1878571 | CNRVC080496 | | Weill et al. Science 2017 |
| **CNRVC030567** | 2003 | Cote d'Ivoire | Africa | ERR1878102 | CNRVC030567 | | Weill et al. Science 2017 |
| **CNRVC030485** | 2003 | Cote d'Ivoire | Africa | ERR1878100 | CNRVC030485 | | Weill et al. Science 2017 |

| | | | | | | | |
|---|---|---|---|---|---|---|---|
| **CNRVC020404** | 2002 | Cote d'Ivoire | Africa | ERR1878089 | CNRVC020404 | | Weill et al. Science 2017 |
| **CNRVC010042** | 2001 | Cote d'Ivoire | Africa | ERR1877647 | CNRVC010042 | | Weill et al. Science 2017 |
| **CNRVC010038** | 2001 | Cote d'Ivoire | Africa | ERR1877646 | CNRVC010038_CAGATC_L001 | | Weill et al. Science 2017 |
| **CNRVC950758** | 1995 | Cote d'Ivoire | Africa | ERR998685 | 16356_8_30 | | Weill et al. Science 2017 |
| **CNRVC950756** | 1995 | Cote d'Ivoire | Africa | ERR998684 | 16356_8_29 | | Weill et al. Science 2017 |
| **CNRVC950755** | 1995 | Cote d'Ivoire | Africa | ERR998683 | 16356_8_28 | | Weill et al. Science 2017 |
| **CNRVC950753** | 1995 | Cote d'Ivoire | Africa | ERR998682 | 16356_8_27 | | Weill et al. Science 2017 |
| **CNRVC980421** | 1998 | Cote d'Ivoire | Africa | ERR976582 | 16244_8_36 | | Weill et al. Science 2017 |
| **CNRVC980420** | 1998 | Cote d'Ivoire | Africa | ERR976581 | 16244_8_35 | | Weill et al. Science 2017 |
| **CNRVC940035** | 1993 | Cote d'Ivoire | Africa | ERR976536 | 16244_7_85 | | Weill et al. Science 2017 |
| **CNRVC940034** | 1994 | Cote d'Ivoire | Africa | ERR976535 | 16244_7_84 | | Weill et al. Science 2017 |
| **CNRVC940030** | 1993 | Cote d'Ivoire | Africa | ERR976534 | 16244_7_83 | | Weill et al. Science 2017 |
| **CNRVC960552** | 1970 | Cote d'Ivoire | Africa | ERR976458 | 16244_7_7 | | Weill et al. Science 2017 |
| **CNRVC910375** | 1991 | Cote d'Ivoire | Africa | ERR976519 | 16244_7_68 | | Weill et al. Science 2017 |
| **CNRVC140141** | 1988 | Cote d'Ivoire | Africa | ERR976512 | 16244_7_61 | | Weill et al. Science 2017 |
| **CNRVC140140** | 1988 | Cote d'Ivoire | Africa | ERR976511 | 16244_7_60 | | Weill et al. Science 2017 |
| **CNRVC950203** | 1970 | Cote d'Ivoire | Africa | ERR976428 | 16244_6_71 | | Weill et al. Science 2017 |
| **CNRVC950191** | 1973 | Cote d'Ivoire | Africa | ERR976427 | 16244_6_70 | | Weill et al. Science 2017 |
| **CNRVC950189** | 1973 | Cote d'Ivoire | Africa | ERR976426 | 16244_6_69 | | Weill et al. Science 2017 |
| **CNRVC950182** | 1973 | Cote d'Ivoire | Africa | ERR976425 | 16244_6_68 | | Weill et al. Science 2017 |
| **CNRVC930171** | 1984 | Cote d'Ivoire | Africa | ERR976411 | 16244_6_54 | | Weill et al. Science 2017 |
| **CNRVC140013** | 2014 | Democratic Republic of the Congo | Africa | ERR1878608 | CNRVC140013 | | Weill et al. Science 2017 |
| **CNRVC140012** | 2014 | Democratic Republic of the Congo | Africa | ERR1878607 | CNRVC140012 | | Weill et al. Science 2017 |

| | | | | | | | |
|---|---|---|---|---|---|---|---|
| **CNRVC080370** | 2008 | Democratic Republic of the Congo | Africa | ERR1878568 | CNRVC080370 | | Weill et al. Science 2017 |
| **CNRVC080133** | 2008 | Democratic Republic of the Congo | Africa | ERR1878567 | CNRVC080133 | | Weill et al. Science 2017 |
| **CNRVC070530** | 2007 | Democratic Republic of the Congo | Africa | ERR1878564 | CNRVC070530 | | Weill et al. Science 2017 |
| **CNRVC070045** | 2007 | Democratic Republic of the Congo | Africa | ERR1878552 | CNRVC070045 | | Weill et al. Science 2017 |
| **CNRVC060153** | 2006 | Democratic Republic of the Congo | Africa | ERR1878155 | CNRVC060153 | | Weill et al. Science 2017 |
| **CNRVC060003** | 2006 | Democratic Republic of the Congo | Africa | ERR1878148 | CNRVC060003 | | Weill et al. Science 2017 |
| **CNRVC050284** | 2005 | Democratic Republic of the Congo | Africa | ERR1878136 | CNRVC050284 | | Weill et al. Science 2017 |
| **CNRVC050077** | 2005 | Democratic Republic of the Congo | Africa | ERR1878132 | CNRVC050077 | | Weill et al. Science 2017 |
| **CNRVC040299** | 2004 | Democratic Republic of the Congo | Africa | ERR1878116 | CNRVC040299 | | Weill et al. Science 2017 |
| **CNRVC030519** | 2002 | Democratic Republic of the Congo | Africa | ERR1878101 | CNRVC030519 | | Weill et al. Science 2017 |
| **CNRVC030469** | 2003 | Democratic Republic of the Congo | Africa | ERR1878097 | CNRVC030469 | | Weill et al. Science 2017 |
| **CNRVC030293** | 2003 | Democratic Republic of the Congo | Africa | ERR1878093 | CNRVC030293 | | Weill et al. Science 2017 |
| **CNRVC030032** | 2003 | Democratic Republic of the Congo | Africa | ERR1878090 | CNRVC030032 | | Weill et al. Science 2017 |
| **CNRVC020284** | 2002 | Democratic Republic of the Congo | Africa | ERR1877955 | CNRVC020284 | | Weill et al. Science 2017 |
| **CNRVC010254** | 2001 | Democratic Republic of the Congo | Africa | ERR1877952 | CNRVC010254 | | Weill et al. Science 2017 |
| **CNRVC010243** | 2001 | Democratic Republic of the Congo | Africa | ERR1877951 | CNRVC010243_GATCAG_L001 | | Weill et al. Science 2017 |

| | | | | | | | |
|---|---|---|---|---|---|---|---|
| **CNRVC970135** | 1997 | Democratic Republic of the Congo | Africa | ERR998746 | 16356_8_91 | | Weill et al. Science 2017 |
| **CNRVC970113** | 1997 | Democratic Republic of the Congo | Africa | ERR998741 | 16356_8_86 | | Weill et al. Science 2017 |
| **CNRVC970064** | 1997 | Democratic Republic of the Congo | Africa | ERR998734 | 16356_8_79 | | Weill et al. Science 2017 |
| **CNRVC970058** | 1997 | Democratic Republic of the Congo | Africa | ERR998733 | 16356_8_78 | | Weill et al. Science 2017 |
| **CNRVC970028** | 1997 | Democratic Republic of the Congo | Africa | ERR998730 | 16356_8_75 | | Weill et al. Science 2017 |
| **CNRVC970027** | 1997 | Democratic Republic of the Congo | Africa | ERR998729 | 16356_8_74 | | Weill et al. Science 2017 |
| **CNRVC970007** | 1997 | Democratic Republic of the Congo | Africa | ERR998724 | 16356_8_69 | | Weill et al. Science 2017 |
| **CNRVC970004** | 1997 | Democratic Republic of the Congo | Africa | ERR998723 | 16356_8_68 | | Weill et al. Science 2017 |
| **CNRVC960308** | 1996 | Democratic Republic of the Congo | Africa | ERR998714 | 16356_8_59 | | Weill et al. Science 2017 |
| **CNRVC960218** | 1996 | Democratic Republic of the Congo | Africa | ERR998698 | 16356_8_43 | | Weill et al. Science 2017 |
| **CNRVC960127** | 1996 | Democratic Republic of the Congo | Africa | ERR998697 | 16356_8_42 | | Weill et al. Science 2017 |
| **CNRVC960124** | 1996 | Democratic Republic of the Congo | Africa | ERR998696 | 16356_8_41 | | Weill et al. Science 2017 |
| **CNRVC940173** | 1994 | Democratic Republic of the Congo | Africa | ERR998659 | 16356_8_4 | | Weill et al. Science 2017 |
| **CNRVC960118** | 1996 | Democratic Republic of the Congo | Africa | ERR998694 | 16356_8_39 | | Weill et al. Science 2017 |
| **CNRVC950746** | 1995 | Democratic Republic of the Congo | Africa | ERR998681 | 16356_8_26 | | Weill et al. Science 2017 |
| **CNRVC950695** | 1995 | Democratic Republic of the Congo | Africa | ERR998669 | 16356_8_14 | | Weill et al. Science 2017 |

| | | | | | | | |
|---|---|---|---|---|---|---|---|
| **CNRVC950694** | 1995 | Democratic Republic of the Congo | Africa | ERR998668 | 16356_8_13 | | Weill et al. Science 2017 |
| **CNRVC950693** | 1995 | Democratic Republic of the Congo | Africa | ERR998667 | 16356_8_12 | | Weill et al. Science 2017 |
| **CNRVC970176** | 1997 | Democratic Republic of the Congo | Africa | ERR976550 | 16244_8_4 | | Weill et al. Science 2017 |
| **CNRVC970174** | 1997 | Democratic Republic of the Congo | Africa | ERR976548 | 16244_8_2 | | Weill et al. Science 2017 |
| **CNRVC940143** | 1994 | Democratic Republic of the Congo | Africa | ERR976544 | 16244_7_93 | | Weill et al. Science 2017 |
| **CNRVC940131** | 1994 | Democratic Republic of the Congo | Africa | ERR976540 | 16244_7_89 | | Weill et al. Science 2017 |
| **CNRVC930185** | 1993 | Democratic Republic of the Congo | Africa | ERR976526 | 16244_7_75 | | Weill et al. Science 2017 |
| **CNRVC930153** | 1993 | Democratic Republic of the Congo | Africa | ERR976525 | 16244_7_74 | | Weill et al. Science 2017 |
| **CNRVC920169** | 1992 | Democratic Republic of the Congo | Africa | ERR976524 | 16244_7_73 | | Weill et al. Science 2017 |
| **CNRVC910429** | 1991 | Democratic Republic of the Congo | Africa | ERR976521 | 16244_7_70 | | Weill et al. Science 2017 |
| **CNRVC980050** | 1984_1986 | Democratic Republic of the Congo | Africa | ERR976487 | 16244_7_36 | | Weill et al. Science 2017 |
| **CNRVC980047** | 1988 | Democratic Republic of the Congo | Africa | ERR976484 | 16244_7_33 | | Weill et al. Science 2017 |
| **CNRVC930170** | 1984 | Democratic Republic of the Congo | Africa | ERR976410 | 16244_6_53 | | Weill et al. Science 2017 |
| **CNRVC930163** | 1988 | Democratic Republic of the Congo | Africa | ERR976404 | 16244_6_47 | | Weill et al. Science 2017 |
| **L384** | 2013 | Democratic Republic of the Congo | Africa | ERR572837 | 12971_7_75 | | Weill et al. Science 2017 |
| **L377** | 2013 | Democratic Republic of the Congo | Africa | ERR572836 | 12971_7_74 | | Weill et al. Science 2017 |

| | | | | | | | |
|---|---|---|---|---|---|---|---|
| **L354** | 2011 | Democratic Republic of the Congo | Africa | ERR572821 | 12971_7_59 | | Weill et al. Science 2017 |
| **L342** | 2001 | Democratic Republic of the Congo | Africa | ERR572810 | 12971_7_48 | | Weill et al. Science 2017 |
| **L339** | 2012 | Democratic Republic of the Congo | Africa | ERR572807 | 12971_7_45 | | Weill et al. Science 2017 |
| **L310** | 2011 | Democratic Republic of the Congo | Africa | ERR572781 | 12971_7_19 | | Weill et al. Science 2017 |
| **L300** | 2011 | Democratic Republic of the Congo | Africa | ERR572772 | 12971_7_10 | | Weill et al. Science 2017 |
| **L284** | 2008 | Democratic Republic of the Congo | Africa | ERR572757 | 12971_6_75 | | Weill et al. Science 2017 |
| **L283** | 2008 | Democratic Republic of the Congo | Africa | ERR572756 | 12971_6_74 | | Weill et al. Science 2017 |
| **L372** | 2013 | Democratic Republic of the Congo | Africa | ERR572563 | 12971_4_50 | | Weill et al. Science 2017 |
| **L363** | 2013 | Democratic Republic of the Congo | Africa | ERR572559 | 12971_4_46 | | Weill et al. Science 2017 |
| **L184** | 2012 | Democratic Republic of the Congo | Africa | ERR572548 | 12971_4_35 | | Weill et al. Science 2017 |
| **L78** | 2009 | Democratic Republic of the Congo | Africa | ERR572538 | 12971_4_25 | | Weill et al. Science 2017 |
| **L1** | 2009 | Democratic Republic of the Congo | Africa | ERR572514 | 12971_4_1 | | Weill et al. Science 2017 |
| **L193** | 2012 | Democratic Republic of the Congo | Africa | ERR386712 | 10868_2_93 | | Weill et al. Science 2017 |
| **L204** | 2012 | Democratic Republic of the Congo | Africa | ERR386711 | 10868_2_92 | | Weill et al. Science 2017 |
| **726** | 2009 | Democratic Republic of the Congo | Africa | ERR386698 | 10868_2_79 | | Weill et al. Science 2017 |
| **L198** | 2012 | Democratic Republic of the Congo | Africa | ERR386695 | 10868_2_76 | | Weill et al. Science 2017 |

| | | | | | | | |
|---|---|---|---|---|---|---|---|
| **3** | 2009 | Democratic Republic of the Congo | Africa | ERR386690 | 10868_2_71 | | Weill et al. Science 2017 |
| **L174** | 2012 | Democratic Republic of the Congo | Africa | ERR386688 | 10868_2_69 | | Weill et al. Science 2017 |
| **2** | 2009 | Democratic Republic of the Congo | Africa | ERR386683 | 10868_2_64 | | Weill et al. Science 2017 |
| **217** | 2009 | Democratic Republic of the Congo | Africa | ERR386672 | 10868_2_53 | | Weill et al. Science 2017 |
| **L201** | 2012 | Democratic Republic of the Congo | Africa | ERR386647 | 10868_2_28 | | Weill et al. Science 2017 |
| **622** | 2009 | Democratic Republic of the Congo | Africa | ERR386629 | 10868_2_10 | | Weill et al. Science 2017 |
| **CNRVC930301** | 1993 | Djibouti | Africa | ERR1879572 | CNRVC930301_GATCAG_L001 | | Weill et al. Science 2017 |
| **CNRVC070020** | 2007 | Djibouti | Africa | ERR1878551 | CNRVC070020 | | Weill et al. Science 2017 |
| **CNRVC010048** | 2000 | Djibouti | Africa | ERR1877938 | CNRVC010048 | | Weill et al. Science 2017 |
| **CNRVC010046** | 2000 | Djibouti | Africa | ERR1877649 | CNRVC010046_CTTGTA_L001 | | Weill et al. Science 2017 |
| **F1393** | 1993 | Djibouti | Africa | ERR1879652 | CDCF1393 | | Weill et al. Science 2017 |
| **CNRVC010018** | 1994 | Djibouti | Africa | ERR976599 | 16244_8_53 | | Weill et al. Science 2017 |
| **CNRVC010017** | 1994 | Djibouti | Africa | ERR976598 | 16244_8_52 | | Weill et al. Science 2017 |
| **CNRVC010015** | 1994 | Djibouti | Africa | ERR976597 | 16244_8_51 | | Weill et al. Science 2017 |
| **CNRVC970177** | 1997 | Djibouti | Africa | ERR976551 | 16244_8_5 | | Weill et al. Science 2017 |
| **CNRVC970175** | 1997 | Djibouti | Africa | ERR976549 | 16244_8_3 | | Weill et al. Science 2017 |
| **CNRVC980158** | 1997 | Djibouti | Africa | ERR976565 | 16244_8_19 | | Weill et al. Science 2017 |
| **CNRVC980044** | 1985 | Djibouti | Africa | ERR976482 | 16244_7_31 | | Weill et al. Science 2017 |
| **CNRVC980039** | 1985 | Djibouti | Africa | ERR976478 | 16244_7_27 | | Weill et al. Science 2017 |
| **CNRVC930027** | 1966 | Egypt | Africa | ERR976397 | 16244_6_40 | | Weill et al. Science 2017 |
| **CNRVC050073** | 2005 | Equatorial Guinea | Africa | ERR1878131 | CNRVC050073 | | Weill et al. Science 2017 |
| **CNRVC050060** | 2005 | Equatorial Guinea | Africa | ERR1878130 | CNRVC050060 | | Weill et al. Science 2017 |

| | | | | | | | |
|---|---|---|---|---|---|---|---|
| **CNRVC150254** | 1998 | Ethiopia | Africa | ERR1879545 | CNRVC150254 | | Weill et al. Science 2017 |
| **CNRVC150218** | 1985 | Ethiopia | Africa | ERR1879392 | CNRVC150218_GTTTCG_L001 | | Weill et al. Science 2017 |
| **CNRVC150213** | 1998 | Ethiopia | Africa | ERR1879387 | CNRVC150213_CCGTCC_L001 | | Weill et al. Science 2017 |
| **CNRVC960596** | 1970 | Ethiopia | Africa | ERR976463 | 16244_7_12 | | Weill et al. Science 2017 |
| **CNRVC930038** | 1970 | Ethiopia | Africa | ERR976399 | 16244_6_42 | | Weill et al. Science 2017 |
| **CNRVC930079** | 1970 | France | Europe | ERR1879569 | CNRVC930079_TCGAAG_L002 | | Weill et al. Science 2017 |
| **CNRVC970170** | 1997 | Gabon | Africa | ERR976547 | 16244_8_1 | | Weill et al. Science 2017 |
| **262_14** | 2014 | Ghana | Africa | ERR1024526 | 16853_4_64 | | Weill et al. Science 2017 |
| **468_14** | 2014 | Ghana | Africa | ERR1024524 | 16853_4_62 | | Weill et al. Science 2017 |
| **19_14** | 2014 | Ghana | Africa | ERR1024521 | 16853_4_59 | | Weill et al. Science 2017 |
| **712_11** | 2011 | Ghana | Africa | ERR1024518 | 16853_4_56 | | Weill et al. Science 2017 |
| **CNRVC960627** | 1970 | Ghana | Africa | ERR976465 | 16244_7_14 | | Weill et al. Science 2017 |
| **CNRVC960623** | 1970 | Ghana | Africa | ERR976464 | 16244_7_13 | | Weill et al. Science 2017 |
| **CNRVC950224** | 1971 | Ghana | Africa | ERR976430 | 16244_6_73 | | Weill et al. Science 2017 |
| **CNRVC950223** | 1971 | Ghana | Africa | ERR976429 | 16244_6_72 | | Weill et al. Science 2017 |
| **CNRVC150217** | 1982 | Ghana/Nigeria | Africa | ERR1879391 | CNRVC150217_GTGGCC_L001 | | Weill et al. Science 2017 |
| **CNRVC900108** | 1990 | Guinea | Africa | ERR1879546 | CNRVC900108_ACAGTG_L001 | | Weill et al. Science 2017 |
| **CNRVC040206** | 2004 | Guinea | Africa | ERR1878112 | CNRVC040206 | | Weill et al. Science 2017 |
| **CNRVC950724** | 1995 | Guinea | Africa | ERR998677 | 16356_8_22 | | Weill et al. Science 2017 |
| **CNRVC950723** | 1995 | Guinea | Africa | ERR998676 | 16356_8_21 | | Weill et al. Science 2017 |
| **CNRVC950699** | 1995 | Guinea | Africa | ERR998670 | 16356_8_15 | | Weill et al. Science 2017 |
| **CNRVC980367** | 1998 | Guinea | Africa | ERR976574 | 16244_8_28 | | Weill et al. Science 2017 |
| **CNRVC940141** | 1994 | Guinea | Africa | ERR976542 | 16244_7_91 | | Weill et al. Science 2017 |
| **CNRVC940128** | 1994 | Guinea | Africa | ERR976539 | 16244_7_88 | | Weill et al. Science 2017 |
| **CNRVC940109** | 1994 | Guinea | Africa | ERR976538 | 16244_7_87 | | Weill et al. Science 2017 |

| | | | | | | | |
|---|---|---|---|---|---|---|---|
| **CNRVC940103** | 1994 | Guinea | Africa | ERR976537 | 16244_7_86 | | Weill et al. Science 2017 |
| **CNRVC960635** | 1970 | Guinea | Africa | ERR976467 | 16244_7_16 | | Weill et al. Science 2017 |
| **CNRVC960629** | 1970 | Guinea | Africa | ERR976466 | 16244_7_15 | | Weill et al. Science 2017 |
| **CNRVC930127** | 1970 | Guinea | Africa | ERR976403 | 16244_6_46 | | Weill et al. Science 2017 |
| **L237** | 2012 | Guinea | Africa | ERR386704 | 10868_2_85 | | Weill et al. Science 2017 |
| **L211** | 2012 | Guinea | Africa | ERR386666 | 10868_2_47 | | Weill et al. Science 2017 |
| **L233** | 2012 | Guinea | Africa | ERR386658 | 10868_2_39 | | Weill et al. Science 2017 |
| **CNRVC120195** | 2012 | Guinea Bissau | Africa | ERR1878605 | CNRVC120195 | | Weill et al. Science 2017 |
| **CNRVC120194** | 2012 | Guinea Bissau | Africa | ERR1878604 | CNRVC120194 | | Weill et al. Science 2017 |
| **K6979** | 2008 | Guinea Bissau | Africa | ERR1880773 | CDCK6979 | | Weill et al. Science 2017 |
| **K6975** | 2008 | Guinea Bissau | Africa | ERR1880772 | CDCK6975 | | Weill et al. Science 2017 |
| **F5144** | 1997 | Guinea Bissau | Africa | ERR1880762 | CDCF5144 | | Weill et al. Science 2017 |
| **CNRVC140133** | 1994-1995 | Guinea Bissau | Africa | ERR976504 | 16244_7_53 | | Weill et al. Science 2017 |
| **CNRVC100278** | 2010 | Haiti | North America | ERR1878592 | CNRVC100278 | | Weill et al. Science 2017 |
| **THSTI_VC63** | 1992 | India | Asia | ERR1880844 | THSTI_VC63 | | Weill et al. Science 2017 |
| **THSTI_VC4** | 1992 | India | Asia | ERR1880843 | THSTI_VC4 | | Weill et al. Science 2017 |
| **THSTI_V8** | 1989 | India | Asia | ERR1880842 | THSTI_V8 | | Weill et al. Science 2017 |
| **THSTI_V157** | 1991 | India | Asia | ERR1880841 | THSTI_V157 | | Weill et al. Science 2017 |
| **THSTI_V156** | 1991 | India | Asia | ERR1880840 | THSTI_V156 | | Weill et al. Science 2017 |
| **THSTI_V149** | 1990 | India | Asia | ERR1880839 | THSTI_V149 | | Weill et al. Science 2017 |
| **THSTI_V12** | 1989 | India | Asia | ERR1880838 | THSTI_V12 | | Weill et al. Science 2017 |
| **THSTI_V108** | 1990 | India | Asia | ERR1880837 | THSTI_V108 | | Weill et al. Science 2017 |
| **THSTI_RC12** | 2001 | India | Asia | ERR1880836 | THSTI_RC12 | | Weill et al. Science 2017 |
| **THSTI_NLC8** | 1999 | India | Asia | ERR1880835 | THSTI_NLC8 | | Weill et al. Science 2017 |
| **THSTI_L9496** | 2006 | India | Asia | ERR1880834 | THSTI_L9496 | | Weill et al. Science 2017 |
| **THSTI_L19089** | 2006 | India | Asia | ERR1880833 | THSTI_L19089 | | Weill et al. Science 2017 |
| **THSTI_K9398** | 2005 | India | Asia | ERR1880832 | THSTI_K9398 | | Weill et al. Science 2017 |

| | | | | | | | |
|---|---|---|---|---|---|---|---|
| **THSTI_K12759** | 2005 | India | Asia | ERR1880831 | THSTI_K12759 | | Weill et al. Science 2017 |
| **THSTI_J4770** | 2004 | India | Asia | ERR1880830 | THSTI_J4770 | | Weill et al. Science 2017 |
| **THSTI_J16173** | 2004 | India | Asia | ERR1880829 | THSTI_J16173 | | Weill et al. Science 2017 |
| **THSTI_IDH02959** | 2010 | India | Asia | ERR1880828 | THSTI_IDH02959 | | Weill et al. Science 2017 |
| **THSTI_IDH02834** | 2010 | India | Asia | ERR1880827 | THSTI_IDH02834 | | Weill et al. Science 2017 |
| **THSTI_IDH02367** | 2009 | India | Asia | ERR1880826 | THSTI_IDH02367 | | Weill et al. Science 2017 |
| **THSTI_IDH01405** | 2009 | India | Asia | ERR1880825 | THSTI_IDH01405 | | Weill et al. Science 2017 |
| **THSTI_IDH01272** | 2008 | India | Asia | ERR1880824 | THSTI_IDH01272 | | Weill et al. Science 2017 |
| **THSTI_IDH00563** | 2008 | India | Asia | ERR1880823 | THSTI_IDH00563 | | Weill et al. Science 2017 |
| **THSTI_IDH00136** | 2007 | India | Asia | ERR1880822 | THSTI_IDH00136 | | Weill et al. Science 2017 |
| **THSTI_IDH0002** | 2007 | India | Asia | ERR1880821 | THSTI_IDH0002 | | Weill et al. Science 2017 |
| **THSTI_I17260** | 2003 | India | Asia | ERR1880820 | THSTI_I17260 | | Weill et al. Science 2017 |
| **THSTI_I16615** | 2003 | India | Asia | ERR1880819 | THSTI_I16615 | | Weill et al. Science 2017 |
| **THSTI_H12381** | 2002 | India | Asia | ERR1880818 | THSTI_H12381 | | Weill et al. Science 2017 |
| **THSTI_H11801** | 2002 | India | Asia | ERR1880817 | THSTI_H11801 | | Weill et al. Science 2017 |
| **THSTI_GP160bis** | 1980 | India | Asia | ERR1880816 | THSTI_GP160bis | | Weill et al. Science 2017 |
| **THSTI_G27875** | 2001 | India | Asia | ERR1880814 | THSTI_G27875 | | Weill et al. Science 2017 |
| **THSTI_E12067** | 1999 | India | Asia | ERR1880813 | THSTI_E12067 | | Weill et al. Science 2017 |
| **THSTI_D31153** | 1998 | India | Asia | ERR1880812 | THSTI_D31153 | | Weill et al. Science 2017 |
| **THSTI_D16133** | 1998 | India | Asia | ERR1880811 | THSTI_D16133 | | Weill et al. Science 2017 |
| **THSTI_CRC198** | 2000 | India | Asia | ERR1880810 | THSTI_CRC198 | | Weill et al. Science 2017 |
| **THSTI_CRC155** | 2000 | India | Asia | ERR1880809 | THSTI_CRC155 | | Weill et al. Science 2017 |
| **THSTI_CO908** | 1995 | India | Asia | ERR1880808 | THSTI_CO908 | | Weill et al. Science 2017 |
| **THSTI_CO783** | 1994 | India | Asia | ERR1880807 | THSTI_CO783 | | Weill et al. Science 2017 |
| **THSTI_CO553** | 1994 | India | Asia | ERR1880806 | THSTI_CO553 | | Weill et al. Science 2017 |

| | | | | | | | |
|---|---|---|---|---|---|---|---|
| **THSTI_CO368** | 1993 | India | Asia | ERR1880805 | THSTI_CO368 | | Weill et al. Science 2017 |
| **THSTI_CO327** | 1993 | India | Asia | ERR1880804 | THSTI_CO327 | | Weill et al. Science 2017 |
| **THSTI_CO1089** | 1995 | India | Asia | ERR1880803 | THSTI_CO1089 | | Weill et al. Science 2017 |
| **THSTI_C18932** | 1997 | India | Asia | ERR1880802 | THSTI_C18932 | | Weill et al. Science 2017 |
| **THSTI_AM305** | 1997 | India | Asia | ERR1880801 | THSTI_AM305 | | Weill et al. Science 2017 |
| **THSTI_AM232** | 1996 | India | Asia | ERR1880800 | THSTI_AM232 | | Weill et al. Science 2017 |
| **THSTI_AM120** | 1996 | India | Asia | ERR1880799 | THSTI_AM120 | | Weill et al. Science 2017 |
| **CNRVC970077** | 1997 | India | Asia | ERR1879637 | CNRVC970077_CCAACA_L002 | | Weill et al. Science 2017 |
| **CNRVC960215** | 1996 | India | Asia | ERR1879597 | CNRVC960215_ATGAGC_L002 | | Weill et al. Science 2017 |
| **CNRVC930066** | 1964 | India | Asia | ERR1879566 | CNRVC930066_ACAGTG_L002 | | Weill et al. Science 2017 |
| **CNRVC930036** | 1962 | India | Asia | ERR1879562 | CNRVC930036_TCGGCA_L001 | | Weill et al. Science 2017 |
| **CNRVC930003** | 1992 | India | Asia | ERR1879553 | CNRVC930003_TCATTC_L001 | | Weill et al. Science 2017 |
| **CNRVC080397** | 2008 | India | Asia | ERR1878570 | CNRVC080397 | | Weill et al. Science 2017 |
| **CNRVC060089** | 2006 | India | Asia | ERR1878151 | CNRVC060089 | | Weill et al. Science 2017 |
| **CNRVC060333** | 2006 | India/Nepal | Asia | ERR1878548 | CNRVC060333 | | Weill et al. Science 2017 |
| **CNRVC010100** | 2001 | Indonesia | Asia | ERR1877943 | CNRVC010100 | | Weill et al. Science 2017 |
| **CNRVC960842** | 1965 | Iran | Asia | ERR1879605 | CNRVC960842_AGTCAA_L001 | | Weill et al. Science 2017 |
| **CNRVC960800** | 1965 | Iran | Asia | ERR1879604 | CNRVC960800_GGCTAC_L001 | | Weill et al. Science 2017 |
| **CNRVC960799** | 1965 | Iran | Asia | ERR1879603 | CNRVC960799_GTGAAA_L001 | | Weill et al. Science 2017 |
| **CNRVC930076** | 1965 | Iran | Asia | ERR1879568 | CNRVC930076_CAGATC_L002 | | Weill et al. Science 2017 |
| **CNRVC960872** | 1965 | Iran | Asia | ERR1879607 | 960872_S21 | | Weill et al. Science 2017 |
| **CNRVC960871** | 1965 | Iran | Asia | ERR1879606 | 960871_S20 | | Weill et al. Science 2017 |
| **CNRVC960768** | 1966 | Iraq | Asia | ERR1879601 | CNRVC960768_TAGCTT_L001 | | Weill et al. Science 2017 |
| **CNRVC950800** | 1995 | Iraq | Asia | ERR1879594 | CNRVC950800_CTATAC_L001 | | Weill et al. Science 2017 |
| **CNRVC950798** | 1995 | Iraq | Asia | ERR1879593 | CNRVC950798_CTAGCT_L001 | | Weill et al. Science 2017 |

| | | | | | | | |
|---|---|---|---|---|---|---|---|
| **CNRVC950792** | 1995 | Iraq | Asia | ERR1879588 | CNRVC950792_GAGTGG_L002 | | Weill et al. Science 2017 |
| **CNRVC030438** | 2003 | Iraq | Asia | ERR1878096 | CNRVC030438_CGGAAT_L001 | | Weill et al. Science 2017 |
| **CNRVC960778** | 1966 | Iraq | Asia | ERR1879602 | 960778_S19 | | Weill et al. Science 2017 |
| **CNRVC960911** | 1970 | Israel | Asia | ERR1879626 | CNRVC960911_AGTTCC_L001 | | Weill et al. Science 2017 |
| **CNRVC960899** | 1970 | Israel | Asia | ERR1879625 | CNRVC960899_CAACTA_L002 | | Weill et al. Science 2017 |
| **CNRVC960891** | 1970 | Israel | Asia | ERR1879624 | 960891_S16 | | Weill et al. Science 2017 |
| **CNRVC930033** | 1970 | Israel | Asia | ERR1879561 | 930033_S15 | | Weill et al. Science 2017 |
| **12_1970** | 1970 | Israel | Asia | ERR042749 | 6353_7_21 | | Weill et al. Science 2017 |
| **4_1970** | 1970 | Israel | Asia | ERR042748 | 6353_7_20 | | Weill et al. Science 2017 |
| **1_1970** | 1970 | Israel | Asia | ERR042745 | 6353_7_17 | | Weill et al. Science 2017 |
| **CNRVC950250** | 1974 | Italy | Europe | ERR1879585 | CNRVC950250_GTTTCG_L002 | | Weill et al. Science 2017 |
| **CNRVC150244** | 1973 | Italy | Europe | ERR1879541 | CNRVC150244_CTCAGA_L001 | | Weill et al. Science 2017 |
| **CNRVC150230** | 1994 | Italy | Europe | ERR1879438 | CNRVC150230_CAGGCG_L001 | | Weill et al. Science 2017 |
| **CNRVC960918** | 1970 | Jordan | Asia | ERR1879627 | 960918_S18 | | Weill et al. Science 2017 |
| **CNRVC150252** | 1998 | Kenya | Africa | ERR1879544 | CNRVC150252 | | Weill et al. Science 2017 |
| **CNRVC150251** | 1998 | Kenya | Africa | ERR1879543 | CNRVC150251 | | Weill et al. Science 2017 |
| **CNRVC150243** | 1998 | Kenya | Africa | ERR1879540 | CNRVC150243_CTATAC_L001 | | Weill et al. Science 2017 |
| **CNRVC150240** | 1998 | Kenya | Africa | ERR1879539 | CNRVC150240_CTAGCT_L001 | | Weill et al. Science 2017 |
| **CNRVC150211** | 1998 | Kenya | Africa | ERR1879385 | CNRVC150211_AGTTCC_L001 | | Weill et al. Science 2017 |
| **CNRVC060120** | 2006 | Kenya | Africa | ERR1878154 | CNRVC060120 | | Weill et al. Science 2017 |
| **CNRVC060119** | 2006 | Kenya | Africa | ERR1878153 | CNRVC060119 | | Weill et al. Science 2017 |
| **CNRVC910416** | 1991 | Kurdistan | Asia | ERR1879550 | CNRVC910416_TAATCG_L001 | | Weill et al. Science 2017 |
| **CNRVC990209** | 1999 | Laos | Asia | ERR1879645 | CNRVC990209_CTAGCT_L002 | | Weill et al. Science 2017 |
| **CNRVC960922** | 1970 | Lebanon | Asia | ERR1879628 | CNRVC960922_CACCGG_L002 | | Weill et al. Science 2017 |
| **CNRVC150223** | 1993 | Lebanon | Asia | ERR1879432 | CNRVC150223_ATGAGC_L001 | | Weill et al. Science 2017 |

| | | | | | | | |
|---|---|---|---|---|---|---|---|
| **CNRVC150215** | 1993 | Lebanon | Asia | ERR1879389 | CNRVC150215_GTCCGC_L001 | | Weill et al. Science 2017 |
| **CNRVC930044** | 1970 | Lebanon | Asia | ERR1879563 | 930044_S17 | | Weill et al. Science 2017 |
| **CNRVC030477** | 2003 | Liberia | Africa | ERR1878099 | CNRVC030477 | | Weill et al. Science 2017 |
| **CNRVC030475** | 2003 | Liberia | Africa | ERR1878098 | CNRVC030475 | | Weill et al. Science 2017 |
| **CNRVC020393** | 2002 | Liberia | Africa | ERR1878088 | CNRVC020393 | | Weill et al. Science 2017 |
| **CNRVC020062** | 2002 | Liberia | Africa | ERR1877954 | CNRVC020062 | | Weill et al. Science 2017 |
| **CNRVC960223** | 1996 | Liberia | Africa | ERR998699 | 16356_8_44 | | Weill et al. Science 2017 |
| **CNRVC940165** | 1994 | Liberia | Africa | ERR998658 | 16356_8_3 | | Weill et al. Science 2017 |
| **CNRVC950730** | 1995 | Liberia | Africa | ERR998679 | 16356_8_24 | | Weill et al. Science 2017 |
| **CNRVC950728** | 1995 | Liberia | Africa | ERR998678 | 16356_8_23 | | Weill et al. Science 2017 |
| **CNRVC940164** | 1994 | Liberia | Africa | ERR998657 | 16356_8_2 | | Weill et al. Science 2017 |
| **CNRVC940162** | 1994 | Liberia | Africa | ERR976546 | 16244_7_95 | | Weill et al. Science 2017 |
| **CNRVC940147** | 1994 | Liberia | Africa | ERR976545 | 16244_7_94 | | Weill et al. Science 2017 |
| **CNRVC940142** | 1994 | Liberia | Africa | ERR976543 | 16244_7_92 | | Weill et al. Science 2017 |
| **CNRVC960946** | 1970 | Liberia | Africa | ERR976469 | 16244_7_18 | | Weill et al. Science 2017 |
| **CNRVC960928** | 1970 | Liberia | Africa | ERR976468 | 16244_7_17 | | Weill et al. Science 2017 |
| **CNRVC960951** | 1970 | Libya | Africa | ERR976471 | 16244_7_20 | | Weill et al. Science 2017 |
| **CNRVC960949** | 1970 | Libya | Africa | ERR976470 | 16244_7_19 | | Weill et al. Science 2017 |
| **CNRVC010055** | 2001 | Madagascar | Africa | ERR1877940 | CNRVC010055_TTAGGC_L001 | | Weill et al. Science 2017 |
| **CNRVC010051** | 2000 | Madagascar | Africa | ERR1877939 | CNRVC010051_ATCACG_L001 | | Weill et al. Science 2017 |
| **CNRVC000065** | 2000 | Madagascar | Africa | ERR976596 | 16244_8_50 | | Weill et al. Science 2017 |
| **CNRVC000064** | 2000 | Madagascar | Africa | ERR976595 | 16244_8_49 | | Weill et al. Science 2017 |
| **CNRVC000062** | 2000 | Madagascar | Africa | ERR976594 | 16244_8_48 | | Weill et al. Science 2017 |
| **CNRVC000060** | 2000 | Madagascar | Africa | ERR976593 | 16244_8_47 | | Weill et al. Science 2017 |
| **CNRVC000059** | 2000 | Madagascar | Africa | ERR976592 | 16244_8_46 | | Weill et al. Science 2017 |

| | | | | | | | |
|---|---|---|---|---|---|---|---|
| **CNRVC000058** | 2000 | Madagascar | Africa | ERR976591 | 16244_8_45 | | Weill et al. Science 2017 |
| **CNRVC000056** | 1999 | Madagascar | Africa | ERR976590 | 16244_8_44 | | Weill et al. Science 2017 |
| **CNRVC000055** | 1999 | Madagascar | Africa | ERR976589 | 16244_8_43 | | Weill et al. Science 2017 |
| **CNRVC990078** | 1999 | Madagascar | Africa | ERR976588 | 16244_8_42 | | Weill et al. Science 2017 |
| **F5097** | 1997 | Malawi | Africa | ERR1880761 | CDCF5097 | | Weill et al. Science 2017 |
| **F5096** | 1997 | Malawi | Africa | ERR1879658 | CDCF5096 | | Weill et al. Science 2017 |
| **CNRVC920049** | 1992 | Malawi | Africa | ERR976523 | 16244_7_72 | | Weill et al. Science 2017 |
| **CNRVC920170** | 1989 | Malawi | Africa | ERR976394 | 16244_6_37 | | Weill et al. Science 2017 |
| **CNRVC900123** | 1990 | Malawi | Africa | ERR976393 | 16244_6_36 | | Weill et al. Science 2017 |
| **CNRVC900117** | 1990 | Malawi | Africa | ERR976392 | 16244_6_35 | | Weill et al. Science 2017 |
| **CNRVC950100** | 1973 | Malaysia | Asia | ERR1879580 | CNRVC950100_CCGTCC_L002 | | Weill et al. Science 2017 |
| **CNRVC060010** | 2005 | Mali | Africa | ERR1878150 | CNRVC060010 | | Weill et al. Science 2017 |
| **CNRVC060009** | 2006 | Mali | Africa | ERR1878149 | CNRVC060009 | | Weill et al. Science 2017 |
| **K0108** | 2003 | Mali | Africa | ERR1880771 | CDCK0108 | | Weill et al. Science 2017 |
| **K0107** | 2003 | Mali | Africa | ERR1880770 | CDCK0107 | | Weill et al. Science 2017 |
| **CNRVC960250** | 1996 | Mali | Africa | ERR998704 | 16356_8_49 | | Weill et al. Science 2017 |
| **CNRVC950708** | 1995 | Mali | Africa | ERR998674 | 16356_8_19 | | Weill et al. Science 2017 |
| **CNRVC950707** | 1995 | Mali | Africa | ERR998673 | 16356_8_18 | | Weill et al. Science 2017 |
| **CNRVC980043** | 1985 | Mali | Africa | ERR976481 | 16244_7_30 | | Weill et al. Science 2017 |
| **CNRVC980041** | 1984 | Mali | Africa | ERR976480 | 16244_7_29 | | Weill et al. Science 2017 |
| **CNRVC960952** | 1970 | Mali | Africa | ERR976472 | 16244_7_21 | | Weill et al. Science 2017 |
| **CNRVC930075** | 1970 | Mali | Africa | ERR976402 | 16244_6_45 | | Weill et al. Science 2017 |
| **CNRVC070144** | 2005 | Mauritania | Africa | ERR1878559 | CNRVC070144 | | Weill et al. Science 2017 |
| **CNRVC070142** | 2006 | Mauritania | Africa | ERR1878558 | CNRVC070142 | | Weill et al. Science 2017 |
| **CNRVC070136** | 2006 | Mauritania | Africa | ERR1878557 | CNRVC070136 | | Weill et al. Science 2017 |

| | | | | | | | |
|---|---|---|---|---|---|---|---|
| **CNRVC070124** | 2005 | Mauritania | Africa | ERR1878556 | CNRVC070124 | | Weill et al. Science 2017 |
| **CNRVC960263** | 1996 | Mauritania | Africa | ERR1879599 | 960263_S14 | | Weill et al. Science 2017 |
| **CNRVC960260** | 1996 | Mauritania | Africa | ERR1879598 | 960260_S13 | | Weill et al. Science 2017 |
| **CNRVC940180** | 1994 | Morocco | Africa | ERR998661 | 16356_8_6 | | Weill et al. Science 2017 |
| **CNRVC940179** | 1994 | Morocco | Africa | ERR998660 | 16356_8_5 | | Weill et al. Science 2017 |
| **CNRVC940137** | 1994 | Morocco | Africa | ERR976541 | 16244_7_90 | | Weill et al. Science 2017 |
| **CNRVC930255** | 1993 | Morocco | Africa | ERR976527 | 16244_7_76 | | Weill et al. Science 2017 |
| **CNRVC910386** | 1991 | Morocco | Africa | ERR976520 | 16244_7_69 | | Weill et al. Science 2017 |
| **CNRVC910349** | 1991 | Morocco | Africa | ERR976518 | 16244_7_67 | | Weill et al. Science 2017 |
| **CNRVC910347** | 1991 | Morocco | Africa | ERR976517 | 16244_7_66 | | Weill et al. Science 2017 |
| **CNRVC950498** | 1972 | Morocco | Africa | ERR976450 | 16244_6_93 | | Weill et al. Science 2017 |
| **CNRVC950487** | 1972 | Morocco | Africa | ERR976449 | 16244_6_92 | | Weill et al. Science 2017 |
| **CNRVC950476** | 1972 | Morocco | Africa | ERR976448 | 16244_6_91 | | Weill et al. Science 2017 |
| **CNRVC950468** | 1972 | Morocco | Africa | ERR976447 | 16244_6_90 | | Weill et al. Science 2017 |
| **CNRVC950466** | 1972 | Morocco | Africa | ERR976446 | 16244_6_89 | | Weill et al. Science 2017 |
| **CNRVC950431** | 1974 | Morocco | Africa | ERR976445 | 16244_6_88 | | Weill et al. Science 2017 |
| **CNRVC900125** | 1990 | Mozambique | Africa | ERR1879547 | CNRVC900125_GCCAAT_L001 | | Weill et al. Science 2017 |
| **F5149** | 1997 | Mozambique | Africa | ERR1880763 | CDCF5149 | | Weill et al. Science 2017 |
| **C8178** | 1992 | Mozambique | Africa | ERR1877616 | CDCC8178_rep | | Weill et al. Science 2017 |
| **B_64** | 2004 | Mozambique | Africa | ERR044797 | 6437_7_20 | | Weill et al. Science 2017 |
| **CNRVC990039** | 1999 | Mozambique | Africa | ERR976587 | 16244_8_41 | | Weill et al. Science 2017 |
| **CNRVC990038** | 1999 | Mozambique | Africa | ERR976586 | 16244_8_40 | | Weill et al. Science 2017 |
| **CNRVC990019** | 1999 | Mozambique | Africa | ERR976585 | 16244_8_39 | | Weill et al. Science 2017 |
| **CNRVC950641** | 1972 | Myanmar | Asia | ERR1879587 | CNRVC950641_GTCCGC_L001 | | Weill et al. Science 2017 |
| **CNRVC930029** | 1971 | Myanmar | Asia | ERR1879559 | CNRVC930029_TCGAAG_L001 | | Weill et al. Science 2017 |

| | | | | | | | |
|---|---|---|---|---|---|---|---|
| **CNRVC940135** | 1994 | Nepal | Asia | ERR1879576 | CNRVC940135_GGCTAC_L002 | | Weill et al. Science 2017 |
| **CNRVC960976** | 1962 | New Guinea | Oceania | ERR1879629 | CNRVC960976_CACGAT_L002 | | Weill et al. Science 2017 |
| **CNRVC140167** | 2010 | Niger | Africa | ERR1878612 | CNRVC140167 | | Weill et al. Science 2017 |
| **CNRVC140160** | 2010 | Niger | Africa | ERR1878611 | CNRVC140160 | | Weill et al. Science 2017 |
| **CNRVC080124** | 2008 | Niger | Africa | ERR1878566 | CNRVC080124 | | Weill et al. Science 2017 |
| **CNRVC080122** | 2008 | Niger | Africa | ERR1878565 | CNRVC080122 | | Weill et al. Science 2017 |
| **CNRVC060273** | 2006 | Niger | Africa | ERR1878545 | CNRVC060273 | | Weill et al. Science 2017 |
| **CNRVC060272** | 2006 | Niger | Africa | ERR1878544 | CNRVC060272 | | Weill et al. Science 2017 |
| **CNRVC040210** | 2004 | Niger | Africa | ERR1878114 | CNRVC040210 | | Weill et al. Science 2017 |
| **CNRVC040209** | 2004 | Niger | Africa | ERR1878113 | CNRVC040209 | | Weill et al. Science 2017 |
| **CNRVC000005** | 1999 | Niger | Africa | ERR1877637 | CNRVC000005_CGATGT_L001 | | Weill et al. Science 2017 |
| **CNRVC940190** | 1994 | Niger | Africa | ERR998663 | 16356_8_8 | | Weill et al. Science 2017 |
| **CNRVC960003** | 1996 | Niger | Africa | ERR998689 | 16356_8_34 | | Weill et al. Science 2017 |
| **CNRVC950759** | 1995 | Niger | Africa | ERR998686 | 16356_8_31 | | Weill et al. Science 2017 |
| **CNRVC910279** | 1991 | Niger | Africa | ERR976516 | 16244_7_65 | | Weill et al. Science 2017 |
| **CNRVC960971** | 1970 | Niger | Africa | ERR976474 | 16244_7_23 | | Weill et al. Science 2017 |
| **CNRVC960959** | 1970 | Niger | Africa | ERR976473 | 16244_7_22 | | Weill et al. Science 2017 |
| **CNRVC140150** | 2014 | Nigeria | Africa | ERR1878610 | CNRVC140150 | | Weill et al. Science 2017 |
| **CNRVC140027** | 2014 | Nigeria | Africa | ERR1878609 | CNRVC140027 | | Weill et al. Science 2017 |
| **CNRVC110262** | 2011 | Nigeria | Africa | ERR1878601 | CNRVC110262 | | Weill et al. Science 2017 |
| **CNRVC110060** | 2011 | Nigeria | Africa | ERR1878594 | CNRVC110060 | | Weill et al. Science 2017 |
| **CNRVC110058** | 2011 | Nigeria | Africa | ERR1878593 | CNRVC110058 | | Weill et al. Science 2017 |
| **CNRVC100224** | 2010 | Nigeria | Africa | ERR1878589 | CNRVC100224 | | Weill et al. Science 2017 |
| **CNRVC100159** | 2010 | Nigeria | Africa | ERR1878583 | CNRVC100159 | | Weill et al. Science 2017 |
| **CNRVC090177** | 2009 | Nigeria | Africa | ERR1878579 | CNRVC090177 | | Weill et al. Science 2017 |

| | | | | | | | |
|---|---|---|---|---|---|---|---|
| **CNRVC090175** | 2009 | Nigeria | Africa | ERR1878578 | CNRVC090175 | | Weill et al. Science 2017 |
| **CNRVC060280** | 2006 | Nigeria | Africa | ERR1878547 | CNRVC060280 | | Weill et al. Science 2017 |
| **CNRVC060279** | 2006 | Nigeria | Africa | ERR1878546 | CNRVC060279 | | Weill et al. Science 2017 |
| **CNRVC050334** | 2005 | Nigeria | Africa | ERR1878147 | CNRVC050334 | | Weill et al. Science 2017 |
| **CNRVC050323** | 2005 | Nigeria | Africa | ERR1878146 | CNRVC050323 | | Weill et al. Science 2017 |
| **F4170** | 1997 | Nigeria | Africa | ERR1879657 | CDCF4170 | | Weill et al. Science 2017 |
| **F4169** | 1997 | Nigeria | Africa | ERR1879656 | CDCF4169 | | Weill et al. Science 2017 |
| **CNRVC960271** | 1996 | Nigeria | Africa | ERR998709 | 16356_8_54 | | Weill et al. Science 2017 |
| **CNRVC960258** | 1996 | Nigeria | Africa | ERR998707 | 16356_8_52 | | Weill et al. Science 2017 |
| **CNRVC960254** | 1996 | Nigeria | Africa | ERR998706 | 16356_8_51 | | Weill et al. Science 2017 |
| **CNRVC960252** | 1996 | Nigeria | Africa | ERR998705 | 16356_8_50 | | Weill et al. Science 2017 |
| **CNRVC960028** | 1996 | Nigeria | Africa | ERR998692 | 16356_8_37 | | Weill et al. Science 2017 |
| **CNRVC950238** | 1970 | Nigeria | Africa | ERR976432 | 16244_6_75 | | Weill et al. Science 2017 |
| **CNRVC950235** | 1970 | Nigeria | Africa | ERR976431 | 16244_6_74 | | Weill et al. Science 2017 |
| **CNRVC961005** | 1964 | Pakistan | Asia | ERR1879630 | CNRVC961005_CACTCA_L002 | | Weill et al. Science 2017 |
| **CNRVC150220** | 1992 | Pakistan | Asia | ERR1879429 | CNRVC150220_GAGTGG_L001 | | Weill et al. Science 2017 |
| **CNRVC090144** | 2009 | Pakistan | Asia | ERR1878577 | CNRVC090144 | | Weill et al. Science 2017 |
| **CNRVC090143** | 2009 | Pakistan | Asia | ERR1878576 | CNRVC090143 | | Weill et al. Science 2017 |
| **CNRVC090123** | 2009 | Pakistan | Asia | ERR1878575 | CNRVC090123 | | Weill et al. Science 2017 |
| **CNRVC050280** | 2005 | Pakistan | Asia | ERR1878135 | CNRVC050280 | | Weill et al. Science 2017 |
| **Gaz11** | 1994 | Palestine | Asia | ERR042733 | 6353_7_5 | | Weill et al. Science 2017 |
| **Gaz21** | 1994 | Palestine | Asia | ERR042738 | 6353_7_10 | | Weill et al. Science 2017 |
| **Gaz2** | 1994 | Palestine | Asia | ERR042729 | 6353_7_1 | | Weill et al. Science 2017 |
| **CNRVC950218** | 1973 | Philippines | Asia | ERR1879584 | CNRVC950218_GTGGCC_L002 | | Weill et al. Science 2017 |
| **CNRVC940044** | 1963 | Philippines | Asia | ERR1879573 | CNRVC940044_ACTTGA_L002 | | Weill et al. Science 2017 |

| | | | | | | | |
|---|---|---|---|---|---|---|---|
| **CNRVC930060** | 1961 | Philippines | Asia | ERR1879564 | CNRVC930060_CGATGT_L002 | | Weill et al. Science 2017 |
| **CNRVC140139** | 1974 | Portugal | Europe | ERR976510 | 16244_7_59 | | Weill et al. Science 2017 |
| **CNRVC140137** | 1974 | Portugal | Europe | ERR976508 | 16244_7_57 | | Weill et al. Science 2017 |
| **CNRVC140136** | 1971 | Portugal | Europe | ERR976507 | 16244_7_56 | | Weill et al. Science 2017 |
| **CNRVC140135** | 1971 | Portugal | Europe | ERR976506 | 16244_7_55 | | Weill et al. Science 2017 |
| **CNRVC140134** | 1971 | Portugal | Europe | ERR976505 | 16244_7_54 | | Weill et al. Science 2017 |
| **NICD_69175** | 1980 | Republic of South Africa | Africa | ERR1880792 | NICD_69175 | | Weill et al. Science 2017 |
| **NICD_69173** | 1980 | Republic of South Africa | Africa | ERR1880791 | NICD_69173 | | Weill et al. Science 2017 |
| **NICD_69147** | 1980 | Republic of South Africa | Africa | ERR1880786 | NICD_69147 | | Weill et al. Science 2017 |
| **NICD_4076** | 2001 | Republic of South Africa | Africa | ERR1880790 | NICD_4076 | | Weill et al. Science 2017 |
| **NICD_273** | 1998 | Republic of South Africa | Africa | ERR1880789 | NICD_273 | | Weill et al. Science 2017 |
| **NICD_16879** | 2001 | Republic of South Africa | Africa | ERR1880788 | NICD_16879 | | Weill et al. Science 2017 |
| **NICD_16756** | 2001 | Republic of South Africa | Africa | ERR1880787 | NICD_16756 | | Weill et al. Science 2017 |
| **NICD_16744** | 2001 | Republic of South Africa | Africa | ERR1877613 | NICD_16744 | | Weill et al. Science 2017 |
| **NICD_16230** | 2001 | Republic of South Africa | Africa | ERR1880785 | NICD_16230 | | Weill et al. Science 2017 |
| **NICD_16211** | 2001 | Republic of South Africa | Africa | ERR1880784 | NICD_16211 | | Weill et al. Science 2017 |
| **NICD_15959** | 2001 | Republic of South Africa | Africa | ERR1880783 | NICD_15959 | | Weill et al. Science 2017 |
| **NICD_15907** | 2001 | Republic of South Africa | Africa | ERR1880782 | NICD_15907 | | Weill et al. Science 2017 |
| **NICD_15104** | 2001 | Republic of South Africa | Africa | ERR1880781 | NICD_15104 | | Weill et al. Science 2017 |
| **NICD_15070** | 2001 | Republic of South Africa | Africa | ERR1880780 | NICD_15070 | | Weill et al. Science 2017 |
| **NICD_14794** | 2001 | Republic of South Africa | Africa | ERR1880779 | NICD_14794 | | Weill et al. Science 2017 |
| **NICD_14191** | 2001 | Republic of South Africa | Africa | ERR1880778 | NICD_14191 | | Weill et al. Science 2017 |
| **NICD_1405** | 2001 | Republic of South Africa | Africa | ERR1880777 | NICD_1405 | | Weill et al. Science 2017 |
| **NICD_1264** | 2001 | Republic of South Africa | Africa | ERR1880776 | NICD_1264 | | Weill et al. Science 2017 |
| **NICD_1242** | 2001 | Republic of South Africa | Africa | ERR1880775 | NICD_1242 | | Weill et al. Science 2017 |

| | | | | | | | |
|---|---|---|---|---|---|---|---|
| **NICD_1241** | 2001 | Republic of South Africa | Africa | ERR1880774 | NICD_1241 | | Weill et al. Science 2017 |
| **F8479** | 2002 | Republic of South Africa | Africa | ERR1880769 | CDCF8479 | | Weill et al. Science 2017 |
| **F8478** | 2002 | Republic of South Africa | Africa | ERR1880768 | CDCF8478 | | Weill et al. Science 2017 |
| **CNRVC990256** | 1999 | Republic of the Congo | Africa | ERR1879647 | CNRVC990256_GTGGCC_L001 | | Weill et al. Science 2017 |
| **CNRVC070012** | 2007 | Republic of the Congo | Africa | ERR1878550 | CNRVC070012 | | Weill et al. Science 2017 |
| **CNRVC070011** | 2007 | Republic of the Congo | Africa | ERR1878549 | CNRVC070011 | | Weill et al. Science 2017 |
| **CNRVC980034** | 1998 | Republic of the Congo | Africa | ERR976560 | 16244_8_14 | | Weill et al. Science 2017 |
| **CNRVC980030** | 1998 | Republic of the Congo | Africa | ERR976559 | 16244_8_13 | | Weill et al. Science 2017 |
| **CNRVC940047** | 1991 | Romania | Europe | ERR1879574 | CNRVC940047_GATCAG_L002 | | Weill et al. Science 2017 |
| **CNRVC150224** | 1990 | Romania | Europe | ERR1879433 | CNRVC150224_ATTCCT_L001 | | Weill et al. Science 2017 |
| **CNRVC150222** | 1987 | Romania | Europe | ERR1879431 | CNRVC150222_ACTGAT_L001 | | Weill et al. Science 2017 |
| **CNRVC150221** | 1991 | Romania | Europe | ERR1879430 | CNRVC150221_GGTAGC_L001 | | Weill et al. Science 2017 |
| **CNRVC150219** | 1994 | Romania | Europe | ERR1879393 | CNRVC150219_CGTACG_L001 | | Weill et al. Science 2017 |
| **CNRVC150214** | 1981 | Romania | Europe | ERR1879388 | CNRVC150214_GTAGAG_L001 | | Weill et al. Science 2017 |
| **CNRVC150134** | 1990 | Russian Federation | Europe | ERR1878616 | 150134_S11 | | Weill et al. Science 2017 |
| **CNRVC150133** | 1990 | Russian Federation | Europe | ERR1878615 | 150133_S10 | | Weill et al. Science 2017 |
| **CNRVC000212** | 2000 | Rwanda | Africa | ERR1877642 | CNRVC000212_ACAGTG_L001 | | Weill et al. Science 2017 |
| **CNRVC000209** | 2000 | Rwanda | Africa | ERR1877641 | CNRVC000209 | | Weill et al. Science 2017 |
| **B9632** | 1998 | Rwanda | Africa | ERR1877615 | CDCB9632_rep | | Weill et al. Science 2017 |
| **B9631** | 1998 | Rwanda | Africa | ERR1877614 | CDCB9631_rep | | Weill et al. Science 2017 |
| **CNRVC960329** | 1996 | Rwanda | Africa | ERR998722 | 16356_8_67 | | Weill et al. Science 2017 |
| **CNRVC960328** | 1996 | Rwanda | Africa | ERR998721 | 16356_8_66 | | Weill et al. Science 2017 |
| **CNRVC960317** | 1996 | Rwanda | Africa | ERR998716 | 16356_8_61 | | Weill et al. Science 2017 |
| **CNRVC960312** | 1996 | Rwanda | Africa | ERR998715 | 16356_8_60 | | Weill et al. Science 2017 |
| **CNRVC960299** | 1996 | Rwanda | Africa | ERR998713 | 16356_8_58 | | Weill et al. Science 2017 |

| | | | | | | | |
|---|---|---|---|---|---|---|---|
| **CNRVC980324** | 1998 | Rwanda | Africa | ERR976569 | 16244_8_23 | | Weill et al. Science 2017 |
| **CNRVC980195** | 1998 | Rwanda | Africa | ERR976567 | 16244_8_21 | | Weill et al. Science 2017 |
| **CNRVC980194** | 1998 | Rwanda | Africa | ERR976566 | 16244_8_20 | | Weill et al. Science 2017 |
| **CNRVC930409** | 1993 | Rwanda | Africa | ERR976528 | 16244_7_77 | | Weill et al. Science 2017 |
| **CNRVC940051** | 1988 | Rwanda | Africa | ERR976414 | 16244_6_57 | | Weill et al. Science 2017 |
| **SaoTome21** | 1989 | Sao Tome | Africa | ERR1880798 | Sao_Tome_21_TGACCA_L001_rep | | Weill et al. Science 2017 |
| **CNRVC920176** | 1989 | Sao Tome | Africa | ERR1879552 | CNRVC920176_ATCACG_L001 | | Weill et al. Science 2017 |
| **CNRVC950340** | 1974 | Senegal | Africa | ERR1879586 | CNRVC950340_CGTACG_L002 | | Weill et al. Science 2017 |
| **CNRVC150226** | 1994 | Senegal | Africa | ERR1879434 | CNRVC150226_CAACTA_L001 | | Weill et al. Science 2017 |
| **CNRVC040314** | 2004 | Senegal | Africa | ERR1878127 | CNRVC040314 | | Weill et al. Science 2017 |
| **CNRVC040303** | 2004 | Senegal | Africa | ERR1878117 | CNRVC040303 | | Weill et al. Science 2017 |
| **CNRVC960327** | 1996 | Senegal | Africa | ERR998720 | 16356_8_65 | | Weill et al. Science 2017 |
| **CNRVC960324** | 1996 | Senegal | Africa | ERR998719 | 16356_8_64 | | Weill et al. Science 2017 |
| **CNRVC960321** | 1996 | Senegal | Africa | ERR998718 | 16356_8_63 | | Weill et al. Science 2017 |
| **CNRVC960318** | 1996 | Senegal | Africa | ERR998717 | 16356_8_62 | | Weill et al. Science 2017 |
| **CNRVC960272** | 1996 | Senegal | Africa | ERR998710 | 16356_8_55 | | Weill et al. Science 2017 |
| **CNRVC960122** | 1996 | Senegal | Africa | ERR998695 | 16356_8_40 | | Weill et al. Science 2017 |
| **CNRVC960031** | 1996 | Senegal | Africa | ERR998693 | 16356_8_38 | | Weill et al. Science 2017 |
| **CNRVC960023** | 1996 | Senegal | Africa | ERR998691 | 16356_8_36 | | Weill et al. Science 2017 |
| **CNRVC960018** | 1996 | Senegal | Africa | ERR998690 | 16356_8_35 | | Weill et al. Science 2017 |
| **CNRVC950345** | 1971 | Senegal | Africa | ERR976436 | 16244_6_79 | | Weill et al. Science 2017 |
| **CNRVC950343** | 1972 | Senegal | Africa | ERR976435 | 16244_6_78 | | Weill et al. Science 2017 |
| **CNRVC950326** | 1974 | Senegal | Africa | ERR976434 | 16244_6_77 | | Weill et al. Science 2017 |
| **CNRVC950253** | 1973 | Senegal | Africa | ERR976433 | 16244_6_76 | | Weill et al. Science 2017 |
| **CNRVC040257** | 2004 | Sierra Leone | Africa | ERR1878115 | CNRVC040257 | | Weill et al. Science 2017 |

| | | | | | | | |
|---|---|---|---|---|---|---|---|
| **CNRVC040203** | 2004 | Sierra Leone | Africa | ERR1878111 | CNRVC040203 | | Weill et al. Science 2017 |
| **CNRVC950712** | 1995 | Sierra Leone | Africa | ERR998675 | 16356_8_20 | | Weill et al. Science 2017 |
| **CNRVC980418** | 1998 | Sierra Leone | Africa | ERR976580 | 16244_8_34 | | Weill et al. Science 2017 |
| **CNRVC980038** | 1986 | Sierra Leone | Africa | ERR976477 | 16244_7_26 | | Weill et al. Science 2017 |
| **CNRVC961014** | 1970 | Sierra Leone | Africa | ERR976475 | 16244_7_24 | | Weill et al. Science 2017 |
| **CNRVC930037** | 1970 | Sierra Leone | Africa | ERR976398 | 16244_6_41 | | Weill et al. Science 2017 |
| **CNRVC961018** | 1970 | Slovakia | Europe | ERR1879632 | 961018_S2 | | Weill et al. Science 2017 |
| **CNRVC961017** | 1970 | Slovakia | Europe | ERR1879631 | 961017_S1 | | Weill et al. Science 2017 |
| **CNRVC930030** | 1970 | Slovakia | Europe | ERR1879560 | 930030_S3 | | Weill et al. Science 2017 |
| **CNRVC940066** | 1994 | Somalia | Africa | ERR1879575 | CNRVC940066_TAGCTT_L002 | | Weill et al. Science 2017 |
| **CNRVC150236** | 1985 | Somalia | Africa | ERR1879537 | CNRVC150236_CCAACA_L001 | | Weill et al. Science 2017 |
| **CNRVC150234** | 1985 | Somalia | Africa | ERR1879536 | CNRVC150234_CATTTT_L001 | | Weill et al. Science 2017 |
| **CNRVC030392** | 2003 | Somalia | Africa | ERR1878095 | CNRVC030392 | | Weill et al. Science 2017 |
| **CNRVC030391** | 2003 | Somalia | Africa | ERR1878094 | CNRVC030391 | | Weill et al. Science 2017 |
| **CNRVC970053** | 1997 | Somalia | Africa | ERR998732 | 16356_8_77 | | Weill et al. Science 2017 |
| **CNRVC990002** | 1999 | Somalia | Africa | ERR976583 | 16244_8_37 | | Weill et al. Science 2017 |
| **CNRVC080383** | 2008 | South Sudan | Africa | ERR1878569 | CNRVC080383 | | Weill et al. Science 2017 |
| **CNRVC070067** | 2007 | South Sudan | Africa | ERR1878554 | CNRVC070067 | | Weill et al. Science 2017 |
| **CNRVC070066** | 2007 | South Sudan | Africa | ERR1878553 | CNRVC070066 | | Weill et al. Science 2017 |
| **RKI-ZBS2-CH8** | 1971 | Spain | Europe | ERR1880796 | RKI-ZBS2-CH8_CTCAGA_L002 | | Weill et al. Science 2017 |
| **CNRVC930174** | 1981 | Sri Lanka | Asia | ERR1879571 | CNRVC930174_CTTGTA_L002 | | Weill et al. Science 2017 |
| **CNRVC960276** | 1996 | Sudan | Africa | ERR1879600 | CNRVC960276_ATTCCT_L002 | | Weill et al. Science 2017 |
| **CNRVC960142** | 1996 | Sudan | Africa | ERR1879595 | CNRVC960142_GGTAGC_L002 | | Weill et al. Science 2017 |
| **CNRVC980384** | 1998 | Sudan | Africa | ERR976575 | 16244_8_29 | | Weill et al. Science 2017 |
| **CNRVC980356** | 1998 | Sudan | Africa | ERR976571 | 16244_8_25 | | Weill et al. Science 2017 |

| | | | | | | | |
|---|---|---|---|---|---|---|---|
| **CNRVC150212** | 1998 | Tanzania | Africa | ERR1879386 | CNRVC150212_ATGTCA_L001 | | Weill et al. Science 2017 |
| **F5333** | 1998 | Tanzania | Africa | ERR1880767 | CDCF5333 | | Weill et al. Science 2017 |
| **F5332** | 1998 | Tanzania | Africa | ERR1880766 | CDCF5332 | | Weill et al. Science 2017 |
| **F3835** | 1997 | Tanzania | Africa | ERR1879655 | CDCF3835 | | Weill et al. Science 2017 |
| **CNRVC980081** | 1998 | Tanzania | Africa | ERR976564 | 16244_8_18 | | Weill et al. Science 2017 |
| **CNRVC980080** | 1998 | Tanzania | Africa | ERR976563 | 16244_8_17 | | Weill et al. Science 2017 |
| **CNRVC930428** | 1993 | Tanzania | Africa | ERR976531 | 16244_7_80 | | Weill et al. Science 2017 |
| **CNRVC961077** | 1963 | Thailand | Asia | ERR1879633 | CNRVC961077exCNRVC961090_TCGAAG_L001 | | Weill et al. Science 2017 |
| **CNRVC930155** | 1993 | Thailand | Asia | ERR1879570 | CNRVC930155_TCGGCA_L002 | | Weill et al. Science 2017 |
| **THSTI_GP106bis** | 1975 | Togo | Africa | ERR1880815 | THSTI_GP106bis | | Weill et al. Science 2017 |
| **CNRVC010108** | 2001 | Togo | Africa | ERR1877944 | CNRVC010108 | | Weill et al. Science 2017 |
| **L409** | 2012 | Togo | Africa | ERR572842 | 12971_7_80 | | Weill et al. Science 2017 |
| **L400** | 2011 | Togo | Africa | ERR572840 | 12971_7_78 | | Weill et al. Science 2017 |
| **L395** | 2010 | Togo | Africa | ERR572839 | 12971_7_77 | | Weill et al. Science 2017 |
| **L413** | 2012 | Togo | Africa | ERR572592 | 12971_4_79 | | Weill et al. Science 2017 |
| **L410** | 2012 | Togo | Africa | ERR572590 | 12971_4_77 | | Weill et al. Science 2017 |
| **L408** | 2011 | Togo | Africa | ERR572589 | 12971_4_76 | | Weill et al. Science 2017 |
| **L403** | 2011 | Togo | Africa | ERR572585 | 12971_4_72 | | Weill et al. Science 2017 |
| **L399** | 2011 | Togo | Africa | ERR572582 | 12971_4_69 | | Weill et al. Science 2017 |
| **L397** | 2010 | Togo | Africa | ERR572580 | 12971_4_67 | | Weill et al. Science 2017 |
| **L390** | 2010 | Togo | Africa | ERR572574 | 12971_4_61 | | Weill et al. Science 2017 |
| **L388** | 2010 | Togo | Africa | ERR572572 | 12971_4_59 | | Weill et al. Science 2017 |
| **L387** | 2010 | Togo | Africa | ERR572571 | 12971_4_58 | | Weill et al. Science 2017 |
| **CNRVC150229** | 1981 | Tunisia | Africa | ERR1879437 | CNRVC150229_CACTCA_L001 | | Weill et al. Science 2017 |
| **CNRVC150228** | 1973 | Tunisia | Africa | ERR1879436 | CNRVC150228_CACGAT_L001 | | Weill et al. Science 2017 |
| **CNRVC950559** | 1971 | Tunisia | Africa | ERR976455 | 16244_7_4 | | Weill et al. Science 2017 |
| **CNRVC950558** | 1971 | Tunisia | Africa | ERR976454 | 16244_7_3 | | Weill et al. Science 2017 |

| | | | | | | | |
|---|---|---|---|---|---|---|---|
| **CNRVC950376** | 1974 | Tunisia | Africa | ERR976441 | 16244_6_84 | | Weill et al. Science 2017 |
| **RKI-ZBS2-CH99** | 1980 | Turkey | Asia | ERR1880797 | RKI-ZBS2-CH99_TAATCG_L002 | | Weill et al. Science 2017 |
| **RKI-ZBS2-CH234** | 1994 | Turkey | Asia | ERR1880795 | RKI-ZBS2-CH234_TCATTC_L002 | | Weill et al. Science 2017 |
| **RKI-ZBS2-CH218** | 1994 | Turkey | Asia | ERR1880794 | RKI-ZBS2-CH218_TATAAT_L002 | | Weill et al. Science 2017 |
| **CNRVC910164** | 1991 | Turkey | Asia | ERR1879549 | CNRVC910164_CTTGTA_L001 | | Weill et al. Science 2017 |
| **CNRVC150216** | 1976 | Turkey | Asia | ERR1879390 | CNRVC150216_GTGAAA_L001 | | Weill et al. Science 2017 |
| **CNRVC000044** | 2000 | Uganda | Africa | ERR1877639 | CNRVC000044_TGACCA_L001 | | Weill et al. Science 2017 |
| **CNRVC000043** | 2000 | Uganda | Africa | ERR1877638 | CNRVC000043 | | Weill et al. Science 2017 |
| **C8799** | 1992 | Uganda | Africa | ERR1877636 | CDCC8799_rep | | Weill et al. Science 2017 |
| **C8798** | 1992 | Uganda | Africa | ERR1877619 | CDCC8798_rep | | Weill et al. Science 2017 |
| **CNRVC980005** | 1998 | Uganda | Africa | ERR976553 | 16244_8_7 | | Weill et al. Science 2017 |
| **CNRVC980004** | 1998 | Uganda | Africa | ERR976552 | 16244_8_6 | | Weill et al. Science 2017 |
| **CNRVC990015** | 1999 | Uganda | Africa | ERR976584 | 16244_8_38 | | Weill et al. Science 2017 |
| **CNRVC150132** | 1974 | Ukraine | Europe | ERR1878614 | 150132_S9 | | Weill et al. Science 2017 |
| **CNRVC150131** | 1970 | Ukraine | Europe | ERR1878613 | 150131_S8 | | Weill et al. Science 2017 |
| **CNRVC920171** | 1983 | Vietnam | Asia | ERR1879551 | CNRVC920171_TACAGC_L001 | | Weill et al. Science 2017 |
| **CNRVC961129** | 1964 | Vietnam | Asia | ERR1879634 | CNRVC961129_CAGGCG_L002 | | Weill et al. Science 2017 |
| **CNRVC950172** | 1969 | Vietnam | Asia | ERR1879583 | CNRVC950172_GTGAAA_L002 | | Weill et al. Science 2017 |
| **CNRVC950165** | 1968 | Vietnam | Asia | ERR1879582 | CNRVC950165_GTCCGC_L002 | | Weill et al. Science 2017 |
| **CNRVC950161** | 1966 | Vietnam | Asia | ERR1879581 | CNRVC950161_GTAGAG_L002 | | Weill et al. Science 2017 |
| **CNRVC100062** | 2010 | Zambia | Africa | ERR1878582 | CNRVC100062 | | Weill et al. Science 2017 |
| **CNRVC100057** | 2010 | Zambia | Africa | ERR1878581 | CNRVC100057 | | Weill et al. Science 2017 |
| **F5198** | 1998 | Zambia | Africa | ERR1880765 | CDCF5198 | | Weill et al. Science 2017 |
| **F5197** | 1998 | Zambia | Africa | ERR1880764 | CDCF5197 | | Weill et al. Science 2017 |
| **336_01** | 2004 | Zambia | Africa | ERR164776 | 8036_3_36 | | Weill et al. Science 2017 |

| | | | | | | | |
|---|---|---|---|---|---|---|---|
| **259_01** | 2004 | Zambia | Africa | ERR164773 | 8036_3_33 | | Weill et al. Science 2017 |
| **329_01** | 2004 | Zambia | Africa | ERR164768 | 8036_3_28 | | Weill et al. Science 2017 |
| **218_02** | 2003 | Zambia | Africa | ERR164766 | 8036_3_26 | | Weill et al. Science 2017 |
| **169_12** | 2003 | Zambia | Africa | ERR164765 | 8036_3_25 | | Weill et al. Science 2017 |
| **20_03** | 1997 | Zambia | Africa | ERR164763 | 8036_3_23 | | Weill et al. Science 2017 |
| **330_02** | 1997 | Zambia | Africa | ERR164760 | 8036_3_20 | | Weill et al. Science 2017 |
| **177_03** | 1997 | Zambia | Africa | ERR164759 | 8036_3_19 | | Weill et al. Science 2017 |
| **329_02** | 1996 | Zambia | Africa | ERR164758 | 8036_3_18 | | Weill et al. Science 2017 |
| **280_02** | 1996 | Zambia | Africa | ERR164757 | 8036_3_17 | | Weill et al. Science 2017 |
| **204_12** | 2003 | Zambia | Africa | ERR044795 | 6437_7_18 | | Weill et al. Science 2017 |
| **354_02** | 1996 | Zambia | Africa | ERR044794 | 6437_7_17 | | Weill et al. Science 2017 |
| **L254** | 2012 | Zambia | Africa | ERR386663 | 10868_2_44 | | Weill et al. Science 2017 |
| **L268** | 2012 | Zambia | Africa | ERR386661 | 10868_2_42 | | Weill et al. Science 2017 |
| **L273** | 2012 | Zambia | Africa | ERR386656 | 10868_2_37 | | Weill et al. Science 2017 |
| **F3403** | 1996 | Zambia_Republic of South Africa | Africa | ERR1879654 | CDCF3403 | | Weill et al. Science 2017 |
| **F3397** | 1996 | Zambia_Republic of South Africa | Africa | ERR1879653 | CDCF3397 | | Weill et al. Science 2017 |
| **Zim_27** | 2009 | Zimbabwe | Africa | ERR044800 | 6437_7_23 | | Weill et al. Science 2017 |
| **Zim_25** | 2009 | Zimbabwe | Africa | ERR044799 | 6437_7_22 | | Weill et al. Science 2017 |
| **Zim_12** | 2009 | Zimbabwe | Africa | ERR044798 | 6437_7_21 | | Weill et al. Science 2017 |
